# Comparative genomics of *Pseudomonas syringae* reveals convergent gene gain and loss associated with specialisation onto cherry (*Prunus avium*)

**DOI:** 10.1101/244715

**Authors:** Michelle T. Hulin, Andrew D. Armitage, Joana G. Vicente, Eric B. Holub, Laura Baxter, Helen J. Bates, John W. Mansfield, Robert W. Jackson, Richard J. Harrison

## Abstract

- Genome-wide analyses of the effector- and toxin-encoding genes were used to examine the phylogenetics and evolution of pathogenicity amongst diverse strains of *Pseudomonas syringae* causing bacterial canker of cherry (*Prunus avium*) including pathovars *P.s* pv. *morsprunorum* (*Psm*) races 1 and 2, *P.s* pv. *syringae* (*Pss*) and *P.s* pv. *avii*.
- Genome-based phylogenetic analyses revealed *Psm* races and *P.s* pv. *avii* clades were distinct and were each monophyletic, whereas cherry-pathogenic strains of *Pss* were interspersed amongst strains from other host species.
- A maximum likelihood approach was used to predict effectors associated with host specialisation on cherry. *Pss* possesses a smaller repertoire of type III effectors but has more toxin biosynthesis clusters compared with *Psm* and *P.s* pv. *avii*. Evolution of cherry pathogenicity was correlated with gain of genes such as *hopAR1* and *hopBB1* through putative phage transfer and horizontal transfer, respectively. By contrast, loss of the *avrPto/hopAB* redundant effector group was observed in cherry-pathogenic clades. Ectopic expression of *hopAB* and *hopC1* triggered the hypersensitive reaction in cherry leaves, confirming computational predictions.
- Cherry canker provides a fascinating example of convergent evolution of pathogenicity that is explained by the mix of effector and toxin repertoires acting on a common host.

## Introduction

*Pseudomonas syringae* is a species complex, associated with plants and the water cycle, comprising several divergent clades that frequently recombine (Young, 2010; Berge *et al*., 2014; Baltrus *et al*., 2017). Currently, thirteen phylogroups, based on Multi-Locus Sequence Typing (MLST), have been described (Parkinson *et al*., 2011). As a plant pathogen, it is globally important, causing disease on over 180 different host species. *P. syringae* is responsible for recurring chronic diseases in perennial crops, such as cherry canker (Lamichhane *et al*., 2014), and also sporadic outbreaks on annual crops, such as bacterial speck of tomato (Şahin, 2001). Individual strains are highly specialised and assigned to over 60 host-specific pathovars; some of these are further divided into races which show host genotype specificity (Joardar *et al*., 2005). This makes *P. syringae* an important model to study the evolution of host specificity (O’Brien *et al*., 2011; Mansfield *et al*., 2012).

High-throughput sequencing has become a major tool in bacterial studies (Edwards & Holt, 2013). With the increasing number of genomes available, population-level studies can now be conducted to ask complex evolutionary questions, such as how disease epidemics emerge and what ecological processes drive the evolution of pathogenicity (Guttman *et al*., 2014; Monteil *et al*., 2016). Before genomic methods were available, mutational studies of *P. syringae* were used to identify functional virulence factors in pathogenesis, such as type III secretion system effector (T3E) repertoires and toxins (Lindgren 1997; Bender *et al*. 1999). Some T3Es were also shown to act as plant defense elicitors or avirulence (*avr*) factors when detected by a corresponding pathogen recognition (R) protein in the host (Jones & Dangl, 2006). *P. syringae* has evolved a functionally redundant repertoire of effectors, which allows pathogen populations to lose/modify expendable *avr* elicitors, with minimal impact on overall virulence (Arnold & Jackson, 2011). It is believed that as pathogen lineages specialise, they fine-tune their effector repertoires to maximize virulence and avoid detection. Host range becomes restricted because the pathogen may lose effectors important for disease on other hosts or gain effectors detected by other plant species (Schulze-Lefert & Panstruga, 2011). Many genomics studies have therefore focused on identifying patterns that link effector complements with particular diseases (Baltrus *et al*. 2011, 2012; O’Brien *et al*. 2012).

Much of the understanding of *P. syringae* – plant interactions has been achieved using herbaceous plant models. Woody, perennial pathosystems provide a greater challenge (Lamichhane *et al*., 2014). Population genomics of *P.s* pv. *actinidiae*, the causal agent of kiwifruit canker, revealed that three pathogen clades, with distinct effector gene sets, have arisen during kiwifruit cultivation (McCann *et al*., 2013, 2017). A study of the olive pathogen *P.s* pv. *savastanoi* revealed that the *hopBL* effector family is over-represented in wood-infecting pathovars (Matas *et al*., 2014). Apart from effectors, genes involved in the metabolism of aromatic compounds, phytohormone production and tolerance to reactive oxygen species have been implicated in virulence on olive (Buonaurio *et al*., 2015). Bartoli *et al*. (2015a) found that the degradation of the aromatic compound catechol was important for symptom development of *P.s* pv. *actinidiae* on kiwifruit. Green *et al*. (2010), identified differences in sucrose metabolism that may dictate the tissue specificity of *P.s* pv. *aesculi* strains that infect horse chestnut. Nowell *et al*. (2016) also identified genes significantly associated with the woody niche. They found candidate effectors, xylose degradation and the α-ketoadipate pathway were associated with this niche.

This study used genomics to examine the evolution of strains that cause bacterial canker on sweet and wild cherry (both *Prunus avium*). Clades of *P. syringae* that constitute the main causal agents of bacterial canker include *P.s* pv. *morsprunorum* (*Psm*) race 1 and race 2 and a *P.s* pv. *syringae* (*Pss*) (Bultreys & Kaluzna, 2010). In addition, *P.s* pv. *avii* causes bacterial canker of wild cherry (Ménard *et al*., 2003). Recently proposed revisions to species names placed *Psm* R1 in *P. amygdali* and *Psm* R2 in *P. avellanae* (Bull *et al*., 2010). As they are part of the *P. syringae* species complex they will be referred to as *Psm* in this study. The cherry-pathogenic clades of *P. syringae* are reported to exhibit differences in virulence, host range and lifestyle (Scortichini 2010; Crosse & Garrett 1966), making the *P. syringae-cherry* pathosystem an intriguing opportunity to study convergent specialisation. The genomic analysis has been coupled with robust pathogenicity testing (Hulin *et al*., 2018) and functional analysis of potential avirulence (avr) genes. This study provides a proof-of-concept that genomics-based predictions can be used to identify candidate genes involved in disease and will likely become the major tool in disease monitoring, diagnostics and host range prediction.

## Materials and methods

### Bacteria, plants and pathogenicity tests

Methods used for bacterial culture and sources of plants were as described in Hulin *et al*. (2018) and are detailed in supplementary methods. Bacterial strains are listed in Table 1. *Escherichia coli* was plated onto Lysogeny Broth Agar (LBA) and grown overnight at 37 °C. Antibiotic concentrations (μg/ml): Kanamycin 50, Gentamycin 10, Spectinomycin 100, Nitrofurantoin 100. X-gal was used at a concentration of 80μg/ml. Tables S1-S3 list the *P. syringae* mutants, plasmids and primers used in this study.

**Table 1:**
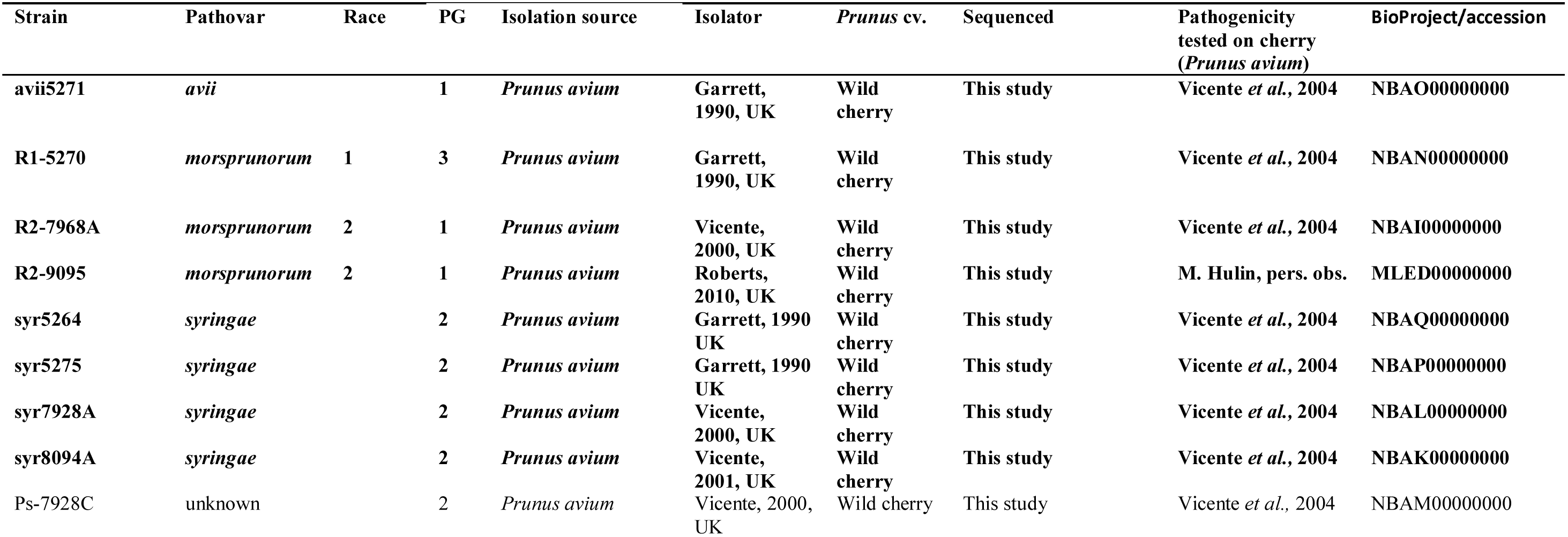
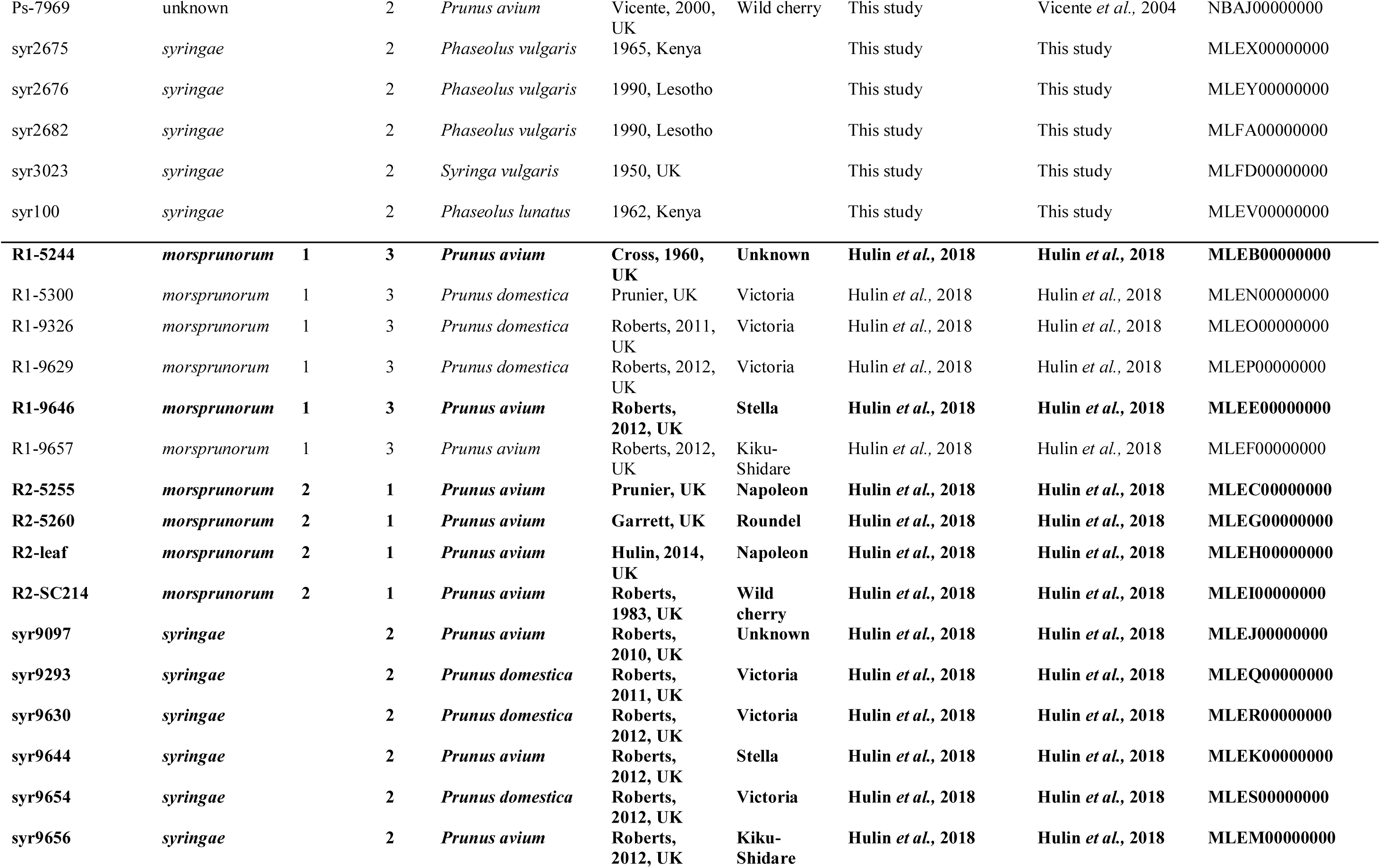
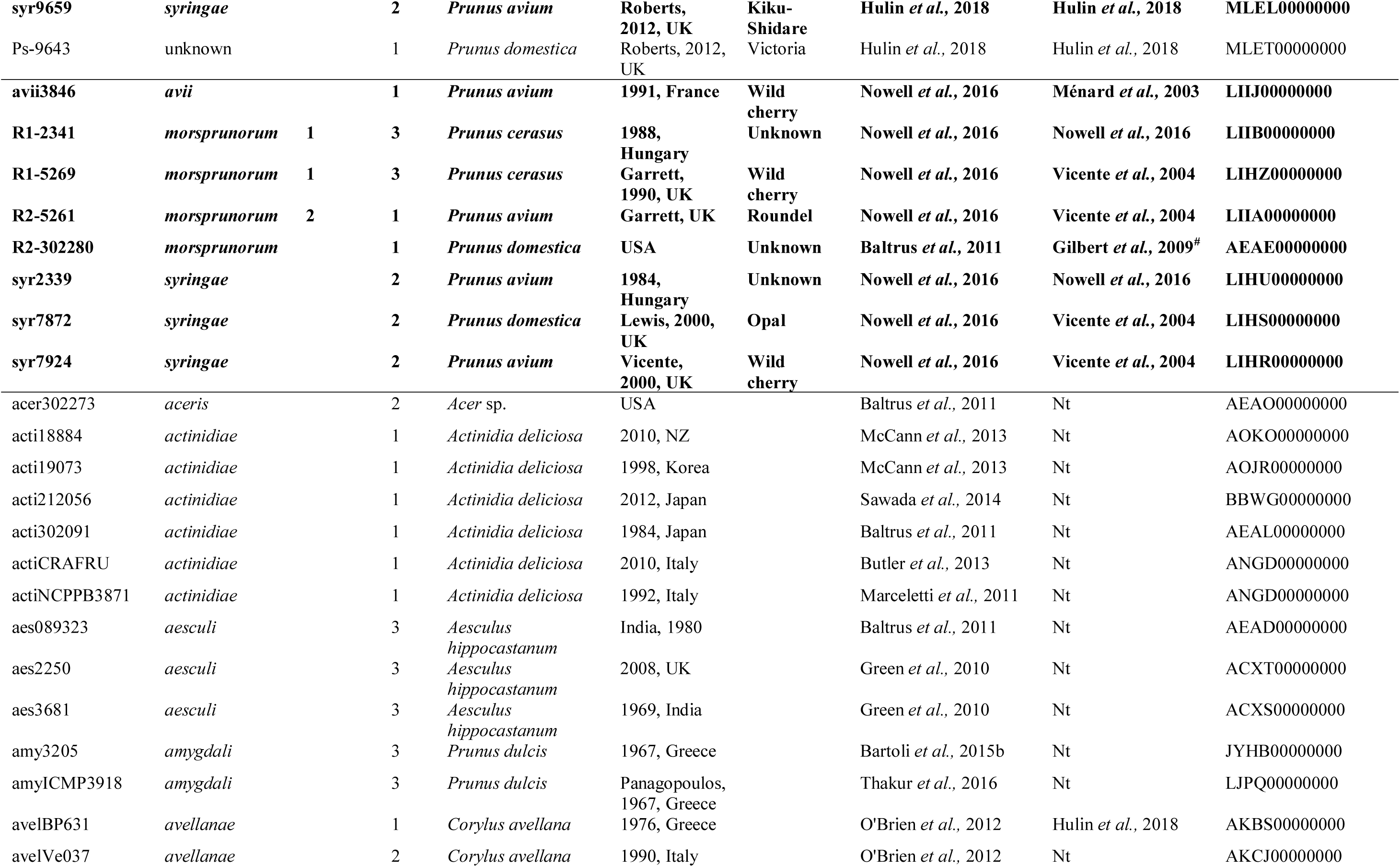
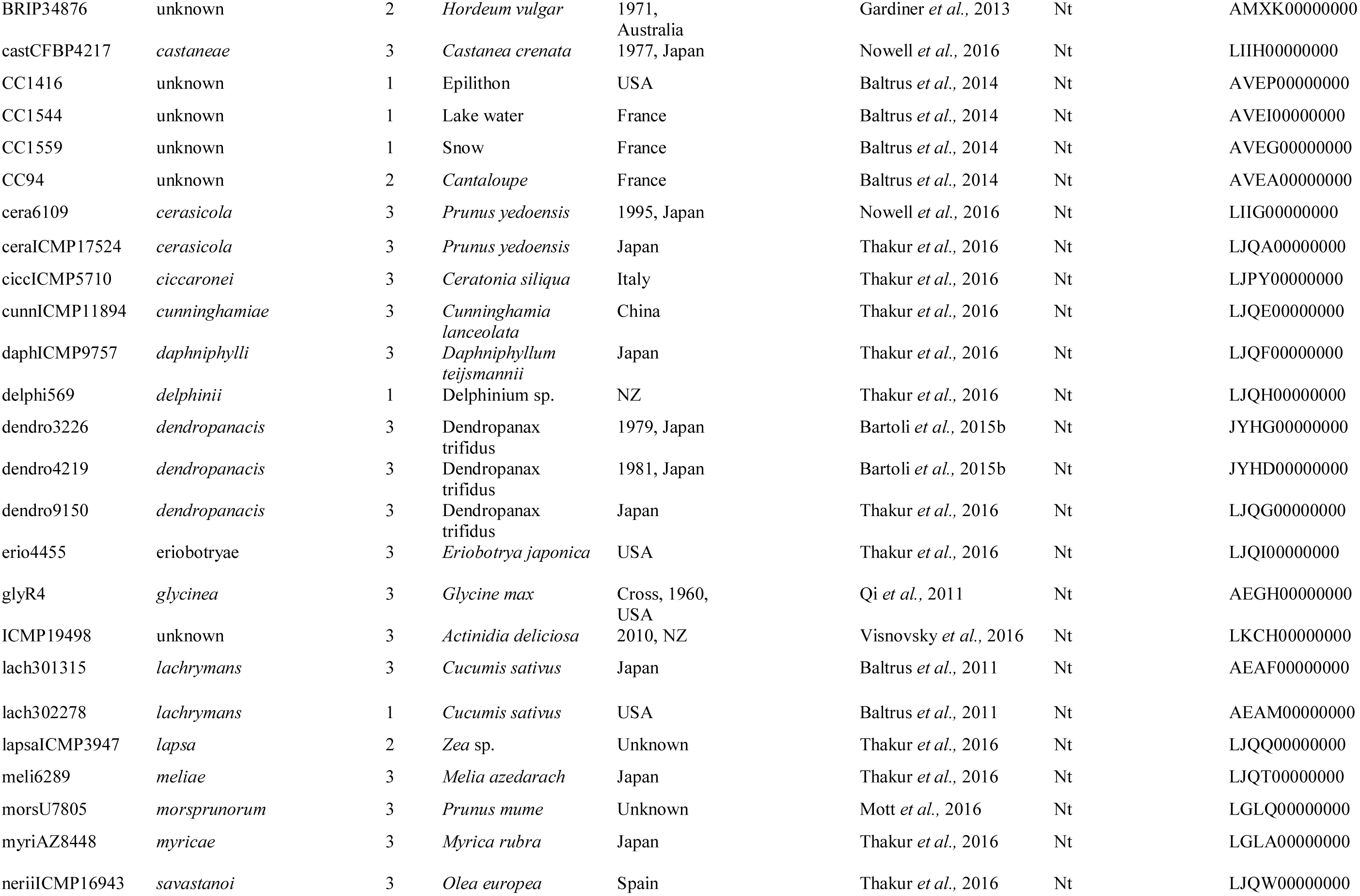
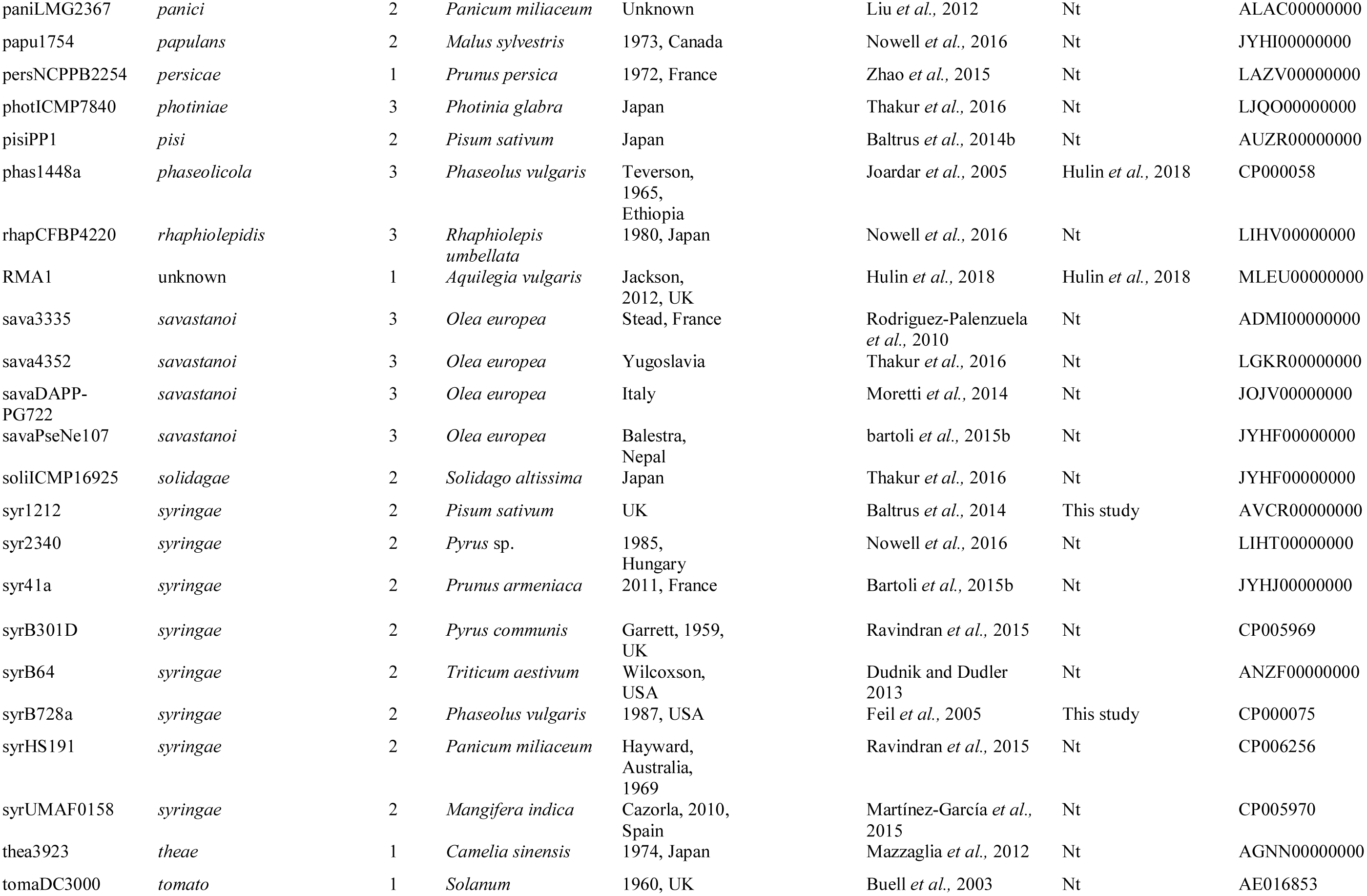
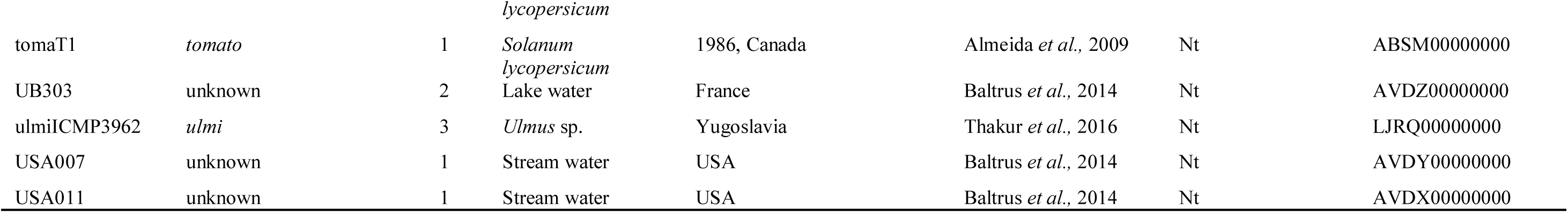
List of bacterial strains used in this study. Pathovar designation, phylogroup, isolation information, cherry pathogenicity (reference for when tested and Nt for not tested) and reference for publication of genome sequence are included. NCBI accession numbers are also listed. Strains in bold were considered pathogenic in cherry. # The pathogenic status of MAFF302280 on cherry is debated. This strain is reported to be the pathotype strain of *P.s* pv. *morsprunorum* (Sawada *et al*., 1999), so is assumed to be equivalent to CFBP 2351, NCPPB2995, ICMP5795 and LMG5075. The strain NCPPB2995 was reported to be potentially non-pathogenic (Gardan *et al*., 1999). In comparison, the ‘same’ strain LMG5075 tested positive for pathogenicity in a recent publication (Gilbert *et al*., 2009). There is no definite link showing that MAFF302280 is the same strain as the others listed as it is not linked to them in online databases (http://www.straininfo.net/) or taxonomy-focused publications (Bull *et al*., 2010). It is assumed to be putatively pathogenic in this study due to its close relatedness to other *Psm* R2 strains, however further pathogenicity tests would be required to fully confirm this.

Pathogenicity tests, performed on detached cherry leaves and analysed as in Hulin *et al*. (2018) are described in the supplementary methods. All ANOVA tables are presented in Tables S17-25.

### Genome sequencing, assembly and annotation

Genome sequencing using Illumina and genome assembly were performed as in Hulin *et al*. (2018). For long-read sequencing, PacBio (Pacific Biosystems) and MinION (Oxford Nanopore) were used. High molecular weight DNA was extracted using a CTAB method (Feil *et al*., 2012). For the PacBio sequencing of *Psm* R1-5244, R2-leaf and syr9097, DNA samples were sent to the Earlham Institute (Norwich) for PacBio RSII sequencing.

For MinION sequencing of *Psm* R1-5300, the DNA library was prepared using the RAD001 rapid-prep kit and run on the Oxford Nanopore MinION, flowcell version R9.5 followed by basecalling using Metrichor. MinION reads were extracted from Fast5 files using Poretools (Loman & Quinlan, 2014). The sequencing reads for both long-read technologies were trimmed and assembled using Canu (Berlin *et al*., 2015) and Circlator was used to circularise contigs (Hunt *et al*., 2015). The assemblies were polished using error-corrected Illumina reads with bowtie2, samtools and Pilon 1.17 (Li *et al*., 2009; Langmead & Salzberg, 2013; Walker *et al*., 2014).

Plasmid profiling was performed using an alkaline-lysis method (Moulton *et al*., 1993) and viewed by gel electrophoresis as in Neale *et al*. (2013). Genomes were submitted to Genbank and accession numbers are listed in Table 1.

### Orthology analysis

OrthoMCL (Li *et al*., 2003) was used to identify orthologous genes. 108 genomes, including those sequenced and those downloaded from NCBI were re-annotated using RAST (Aziz *et al*., 2008) to ensure similar annotation quality. For this reason, the Illumina short-read assemblies of the four long-read sequenced genomes (R1-5244, R1-5300, R2-leaf and syr9097) were used in orthology analysis. OrthoMCL was run with default settings and a 50 residue cut-off length.

### Phylogenetic and genomic analysis of *P. syringae*

Single-copy genes present in all genomes were aligned using ClustalW (Larkin *et al*., 2007) and trimmed using Gblocks (Castresana, 2000). Gene alignments were concatenated using Geneious (Kearse *et al*., 2012). The program jmodeltest 2.1.10 (Posada, 2008) determined the correct evolutionary model for each gene. RAxML-AVX v8.1.15 (Stamatakis, 2014) was used in partitioned mode to build the maximum likelihood phylogeny, with a GTR gamma model and 100 non-parametric bootstrap replicates. To detect core genes that may have undergone recombination, the program GENECONV (Sawyer, 1989) was used as in Yu *et al*. (2016). Whole genome alignments were performed using progressiveMauve (Darling *et al*., 2010).

### Virulence and mobility gene identification

All T3E encoding sequences were downloaded from pseudomonas-syringae.org, including the recent classification of HopF effectors into four alleles (Lo *et al*., 2016). tBLASTn (Altschul *et al*., 1990), was used to search for homologues with a score of ≥70% identity and ≥40% query length to at least one sequence in each effector family. Nucleotide sequences were manually examined for frameshifts or truncations. Disrupted genes were classed as pseudogenes. A heatmap of effector presence was generated using heatmap.2 in gplots (Warnes *et al*., 2016). Interproscan (Quevillon *et al*., 2005) identified protein domains and Illustrator for Biological Sequences (IBS) was used for visualisation (Liu *et al*., 2015). Genomic regions containing effectors were aligned using MAFFT (Katoh *et al*., 2002).

A similar analysis was performed for phytotoxin biosynthesis, wood-degradation, ice nucleation and plasmid-associated genes. Protein sequences were obtained from NCBI (Table S4) and blasted against the genome sequence as above. Prophage identification was performed using PHASTER (Arndt *et al*., 2016).

### Gain and loss analysis

GLOOME (Gain Loss Mapping Engine) was used to plot the gain and loss of genes on the core genome phylogeny (Cohen *et al*., 2010). Effector genes were considered present even if predicted to be pseudogenes, as these can still be gained and lost. The optimisation level was set to “very high”, a mixture model allowing variable gain and loss distributions was used and the distribution type was GENERAL_GAMMA_PLUS_INV. Highly probable events (probability ≥0.80) on the branches leading to cherry-pathogenic strains were extracted.

### BayesTraits analysis

BayesTraits was used to correlate T3E gene evolution with pathogenicity (Pagel, 2004). A binary matrix was created of effector family presence and pathogenicity of each strain. The effector matrix was collapsed into effector families as the different alleles were predicted to perform similar biological functions *in planta* (Cunnac *et al*., 2011). Putative pseudogenes were considered absent as they may be non-functional. A full description of the BayesTraits methodology is described in the supplementary methods.

### Horizontal gene transfer analysis

For each effector family, best hit nucleotide sequences were aligned using clustalW (Larkin *et al*., 2007). RAxML was used to build a phylogenetic tree with a GTR model of evolution and 1000 bootstrap replicates. Incongruence with the core genome tree was examined visually. To further assess horizontal transfer, a species-gene tree reconciliation method RANGER-DTL (Bansal *et al*., 2012) was used, as in Bruns *et al*. (2017). Full details of this are described in the supplementary methods.

### Identification of genomic islands

Genomic islands (GIs) were identified in the PacBio-sequenced strains using IslandViewer3 (Dhillon *et al*., 2015). Islands were manually delimited as in McCann *et al*. (2013). BLASTn was utilised to determine if these GIs were present in other *P. syringae* strains. As most GIs exceeded 10kb, the islands were split into 0.5kb sections prior to analysis. An island was concluded to be fully present if all sections produced hits with a query length >30%. To validate this approach, the Illumina-sequenced genome assemblies of the PacBio-sequenced strains were searched for their own islands.

### General DNA manipulations and bacterial transformations

Cloning and other molecular biology techniques including ectopic expression of potential *avr* genes were as described in earlier works (Staskawicz *et al*., 1984; Arnold *et al*., 2001; Kvitko & Collmer, 2011). Details are provided in the supplementary methods.

## Results

### Genome assembly and sequencing statistics

Genome information gathered in this study enabled a comprehensive analysis and meaningful comparisons to investigate the evolution of pathogenicity amongst *P. syringae* pathogens of cherry. Eighteen *P. syringae* strains isolated from cherry and plum were phenotyped for pathogenicity and genome sequenced in a previous study (Hulin *et al*., 2018). To increase this sample, nine strains isolated from wild cherry and five additional *non-Prunus* out-groups were genome-sequenced using the Illumina MiSeq. The genomes of eight cherry strains sequenced elsewhere were also downloaded from NCBI (Baltrus *et al*., 2011; Nowell *et al*., 2016).

Information on the origin and pathogenicity of each strain is summarised in Table 1. Twenty-eight were considered pathogenic to cherry including all *Pss* isolated from cherry and plum. In contrast, three *Psm* race 1 strains from plum (R1-5300, R1-9326 and R1-9629) and one from cherry strain (R1-9657) failed to induce canker on cherry following tree inoculations; and three strains of unknown taxonomy isolated from plum and cherry (Ps-9643, Ps-7928C and Ps-7969) were non-pathogenic (references in Table 1). The cherry pathogens are referred to as their described pathovar names throughout this study. To simplify figures, the first few letters of the pathovar name were used. “Pss” becomes “syr”, as otherwise *Pss* could refer to other pathovars beginning with “s”, e.g. *savastanoi*.

All cherry/plum isolated strains included in this study were sequenced using Illumina MiSeq. Three representative strains (R1-5244, R2-leaf and syr9097) were sequenced using PacBio and the non-pathogenic *Psm* R1 strain R1-5300 was sequenced using Oxford Nanopore, to obtain more complete genomes. Table 2 summarises the genome assembly statistics for all strains sequenced in this study and Hulin *et al*. (2018). Illumina genomes assembled into 23-352 contigs, whilst the long-read sequenced genomes assembled into 1-6 contigs. The numbers of plasmids in each strain was determined by plasmid profiling (Fig. S1). *Psm* R1 and R2 strains possessed between 2-6 plasmids, *P.s*. pv. *avii* 5271 possessed six plasmids, whereas, apart from three strains (syr5275, syr7928A, syr9644) with one plasmid each, most cherry-pathogenic strains of *Pss* did not possess plasmids. The strain syr9097 that was PacBio-sequenced lacked plasmids. The long-read sequenced genomes all assembled into the correct number of contigs corresponding to chromosome and plasmids, apart from R1-5300. The chromosome of this strain was separated into two contigs (tig0 and tig75) based on a whole genome alignment with *Psm* R1 strain R1-5244 (Fig. S2).

**Table 2:** Assembly statistics for all strains in this study and by Hulin *et al*. (2018). Cherry pathogens are in bold. N50: The weighted median contig size in the assembly. Features: The number of protein encoding and RNA sequences in the annotated genome.

| Assembly | Sequencing | Reference | No. contigs | Plasmids | Total length | GC % | N50 | Average coverage | Features |
| --- | --- | --- | --- | --- | --- | --- | --- | --- | --- |
| **R1-5270** | **Illumina** | **This study** | **185** | **3** | **6258313** | **58.10** | **202152** | **134** | **5770** |
| **R1-5244** | **Illumina** | **Hulin *et al.*, 2018** | **198** | **4** | **6302385** | **58.08** | **227422** | **265** | **5810** |
| R1-5300 | Illumina | Hulin *et al.*, 2018 | 201 | 4 | 6342586 | 57.88 | 142021 | 180 | 5844 |
| R1-9326 | Illumina | Hulin *et al.*, 2018 | 268 | 4 | 6353636 | 57.91 | 142021 | 81 | 5874 |
| R1-9629 | Illumina | Hulin *et al.*, 2018 | 216 | 3 | 6341664 | 57.94 | 142021 | 172 | 5856 |
| **R1-9646** | **Illumina** | **Hulin *et al.*, 2018** | **171** | **3** | **6302776** | **58.03** | **235429** | **180** | **5801** |
| R1-9657 | Illumina | Hulin *et al.*, 2018 | 191 | 4 | 6317852 | 57.91 | 145272 | 158 | 5848 |
| **R2-5255** | **Illumina** | **Hulin *et al.*, 2018** | **206** | **2** | **6448834** | **58.38** | **102760** | **112** | **5966** |
| **R2-7968A** | **Illumina** | **This study** | **278** | **6** | **6498711** | **58.42** | **91262** | **134** | **6016** |
| **R2-5260** | **Illumina** | **Hulin *et al.*, 2018** | **223** | **3** | **6495620** | **58.41** | **101794** | **458** | **5995** |
| **R2-9095** | **Illumina** | **This study** | **304** | **2** | **6418849** | **58.48** | **92453** | **100** | **5887** |
| **R2-leaf** | **Illumina** | **Hulin *et al.*, 2018** | **203** | **5** | **6366714** | **58.48** | **100658** | **180** | **5846** |
| **R2-SC214** | **Illumina** | **Hulin *et al.*, 2018** | **203** | **3** | **6253818** | **58.56** | **108341** | **180** | **5747** |
| **avii5271** | **Illumina** | **This study** | **352** | **6** | **6243644** | **58.56** | **56064** | **127** | **5809** |
| Ps-9643 | Illumina | Hulin *et al.*, 2018 | 58 | 1 | 5937102 | 58.78 | 243355 | 212 | 5386 |
| **syr9097** | **Illumina** | **Hulin *et al.*, 2018** | **66** | **0** | **5892389** | **59.35** | **316078** | **158** | **5117** |
| **syr9293** | **Illumina** | **Hulin *et al.*, 2018** | **73** | **0** | **6135031** | **58.84** | **557853** | **196** | **5302** |
| **syr9630** | **Illumina** | **Hulin *et al.*, 2018** | **57** | **0** | **5940819** | **59.33** | **347701** | **206** | **5175** |
| **syr9644** | **Illumina** | **Hulin *et al.*, 2018** | **75** | **1** | **6173193** | **59.13** | **251053** | **208** | **5334** |
| **syr9654** | **Illumina** | **Hulin *et al.*, 2018** | **49** | **0** | **5941610** | **59.37** | **245023** | **147** | **5148** |
| **syr9656** | **Illumina** | **Hulin *et al.*, 2018** | **39** | **0** | **5980728** | **59.10** | **1007808** | **205** | **5184** |
| **syr9659** | **Illumina** | **Hulin *et al.*, 2018** | **51** | **0** | **5943090** | **59.37** | **235830** | **116** | **5148** |
| **syr5264** | **Illumina** | **This study** | **59** | **0** | **6029896** | **59.08** | **380149** | **114** | **5314** |
| **syr5275** | **Illumina** | **This study** | **64** | **1** | **5994091** | **59.30** | **371492** | **145** | **5207** |
| **syr7928A** | **Illumina** | **This study** | **59** | **1** | **6129363** | **59.26** | **371492** | **141** | **5338** |
| **syr8094A** | **Illumina** | **This study** | **71** | **0** | **5942438** | **59.33** | **265238** | **106** | **5184** |
| syr7928C | Illumina | This study | 49 | 1 | 5994455 | 59.17 | 325175 | 124 | 5318 |
| syr7969 | Illumina | This study | 92 | 0 | 6185932 | 59.01 | 164374 | 151 | 5476 |
| RMA1 | Illumina | Hulin *et al.*, 2018 | 95 | 1 | 6306889 | 58.73 | 187448 | 320 | 5825 |
| syr100 | Illumina | This study | 23 | 0 | 5872916 | 59.33 | 893822 | 83 | 5140 |
| syr2675 | Illumina | This study | 65 | 0 | 5994384 | 59.34 | 227612 | 83 | 5177 |
| syr2676 | Illumina | This study | 90 | 1 | 6158476 | 59.30 | 259660 | 78 | 5387 |
| syr2682 | Illumina | This study | 185 | 1 | 6259099 | 59.21 | 242212 | 84 | 5405 |
| syr3023 | Illumina | This study | 228 | 0 | 6203212 | 58.90 | 456738 | 88 | 5365 |
| **R1-5244** | **PacBio** | **This study** | **5** | **4** | **6445963** | **58.05** | **6109228** | **82** | **6024** |
| **R2-leaf** | **PacBio** | **This study** | **6** | **5** | **6576340** | **58.41** | **6242845** | **141** | **6093** |
| **syr9097** | **PacBio** | **This study** | **1** | **0** | **5929959** | **59.30** | **5929959** | **100** | **5147** |
| R1-5300 | MinION | This study | 6 | 4 | 6645601 | 57.87 | 5688034 | 100 | 6449 |

The *Psm* R1 (R1-5244, R1-5300) and *Psm* R2 (R2-leaf) long-read assemblies contained putatively complete plasmid contigs. These were confirmed to contain plasmid-associated genes (Tables S5-S7). All three strains (R1-5244, R1-5300 and R2-leaf) possessed plasmids with *repA* homologues, indicating they may belong to common plasmid family pPT23A (Zhao *et al*., 2005). Several plasmids also contained conjugational machinery so may be conjugative.

### Core-genome phylogenetic analysis

To examine the relatedness of strains, an analysis of core genes was carried out. 108 genomes of strains from the well studied phylogroups 1-3 isolated from both plants and aquatic environments were selected. A maximum likelihood phylogeny based on 1035 concatenated core genes was constructed using RAxML (Fig. S3). There was low support for certain P2 and P3 clades based on bootstrap analysis. To determine if particular taxa were causing low support, the analysis was systematically repeated for the two phylogroups, with non-cherry strains removed. Support and tree likelihood values were compared (Table S8). Within P3, the removal of *P.s* pv. *eriobotryae* or *P.s* pv. *daphniphylli* improved support, whilst the removal of *P.s* pv. *syringae* 1212 improved support values in P2 (Fig. S4 and S5). The global analysis was then repeated with these taxa removed (Fig. S6-S9). The final phylogeny (Fig. 1), with the highest mean branch support (92.8%) lacked *P.s*. pv. *eriobotryae*. The phylogeny, built using a 611,888bp alignment, contained 102 taxa due to the removal of identical strains (dendro4219, syr9630, R1-9629, R1-9326 and R1-5269). Most support values exceeded 70%, with good support for branches leading to cherry-pathogenic clades.

**Fig. 1:**
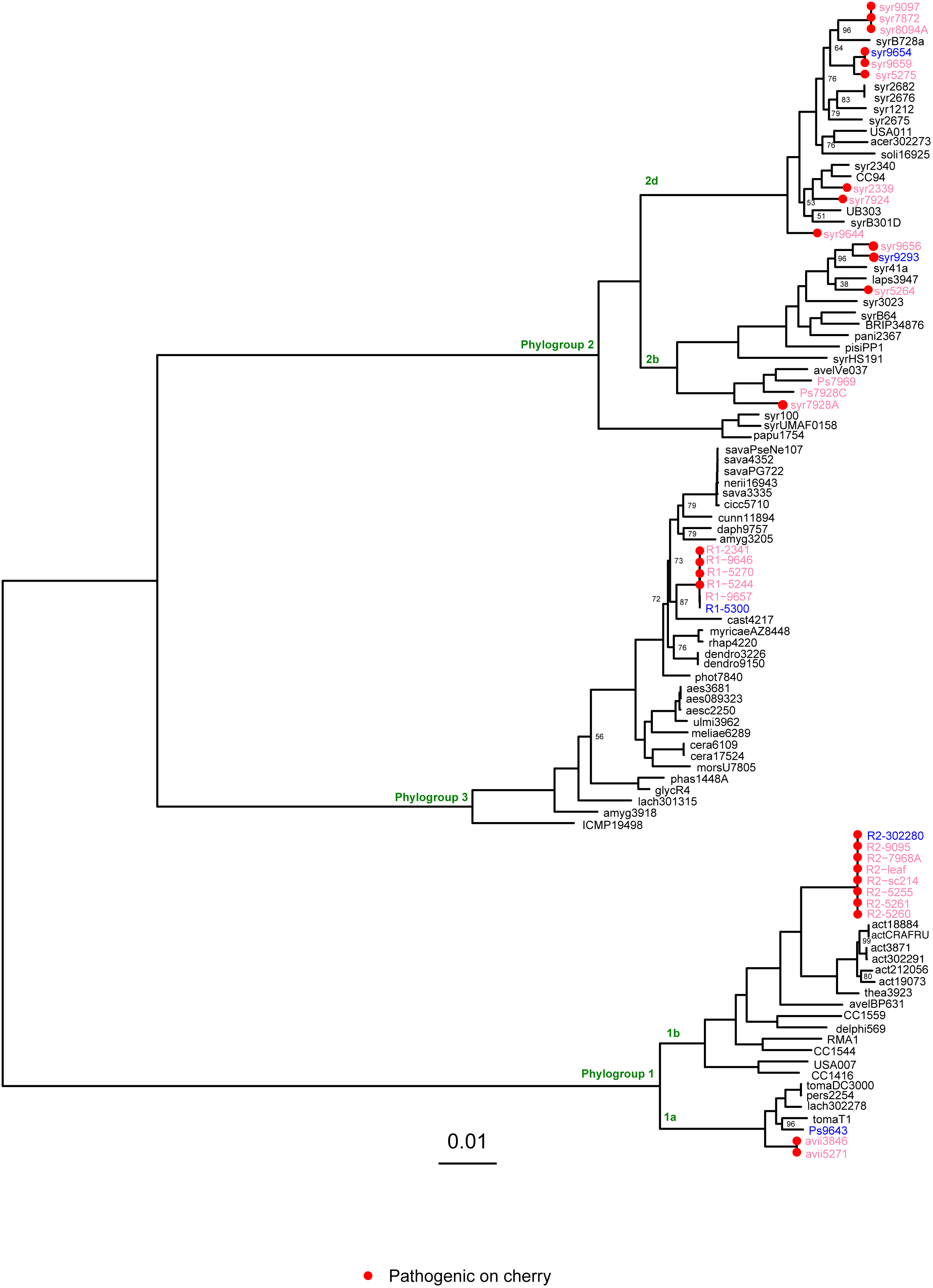
Core genome phylogenetic tree. Multi-locus phylogeny based on 1035 genes which represent the core genome of *P. syringae*. Strains from cherry and plum are highlighted in pink and blue respectively. Strains pathogenic to cherry (assessed in Hulin *et al*., 2018 and Vicente *et al*., 2004) are labelled with red circles. * indicates non-pathogenic to cherry in controlled pathogenicity tests. Phylogroups are also labelled for reference. Percentage bootstrap support values below 99% are shown for each node. The scale is nucleotide substitutions per site.

One explanation for the low support within P2 and P3 was that these clades have undergone core genome recombination. The program GENECONV (Sawyer, 1989) showed that 140 genes had putatively recombined (127,288bp total length, 20.8% of the alignment). Table S9 lists the number of recombination events per phylogroup. It was observed that the most frequent core gene recombination occurred in P3 (73 genes affected), followed by 31 genes in P2 and only 13 in P1.

Cherry pathogens were found in all three phylogroups. The two *Psm* races (R1 in P3, R2 in P1) and *P.s* pv. *avii* (P1) formed monophyletic clades. Within *Psm* R1, cherry pathogenic strains formed a distinct clade from previously classified non-pathogenic strains (Hulin *et al*., 2018). This indicated there has been divergence in their core genomes. By contrast, *Prunus*-infecting strains of *Pss* were found across P2, interspersed with strains isolated from other plants and aquatic environments. To ensure that genomic comparisons between P2 strains were based on differential pathogenicity, several closely related *non-Prunus* strains were pathogenicity tested on detached cherry leaves (Fig. S10). *In planta* bacterial populations of *non-Prunus* strains were reduced compared to two cherry and plum *Pss* strains, indicating that host specificity may exist in P2.

## Search for virulence factors

### The *hrp* pathogenicity island

All sequenced strains were confirmed to contain the *hrp* pathogenicity island required for conventional Type III secretion. Core effectors genes from the conserved effector locus (CEL, Alfano *et al*., 2000) such as *avrE1, hopM1* and *hopAA1*, were also present (Fig. S11), However, *hopAA1* was truncated in both *Psm* R1 and R2 due to inversion events. The *hopAA1* gene was truncated in *Psm* R2, whilst in *Psm* R1 both *hopAA1* and *hopM1* were truncated (Fig. S12).

### Type III effectors and other virulence genes

All 102 genomes used in the phylogenetic analysis were scanned for known T3Es and non-T3 virulence factors. A heatmap of virulence factor presence, absence and pseudogenisation was constructed (Fig. 2). In terms of T3Es, there was variation both between and within the different cherry-pathogenic clades. Notably, *Psm* R1 which contained pathogenic and non-pathogenic strains on cherry showed clear differentiation in effector repertoire (Table S10). *Psm* R1, R2 and *P.s* pv. *avii* possessed 24-34 effector genes, whereas *Pss* strains possessed 9-15. This reduced effector repertoire of *Pss* was representative of P2 strains as noted by Dudnik & Dudler (2014). Table 3 lists the effectors in each long-read genome assembly in order of appearance.

**Fig. 2:**
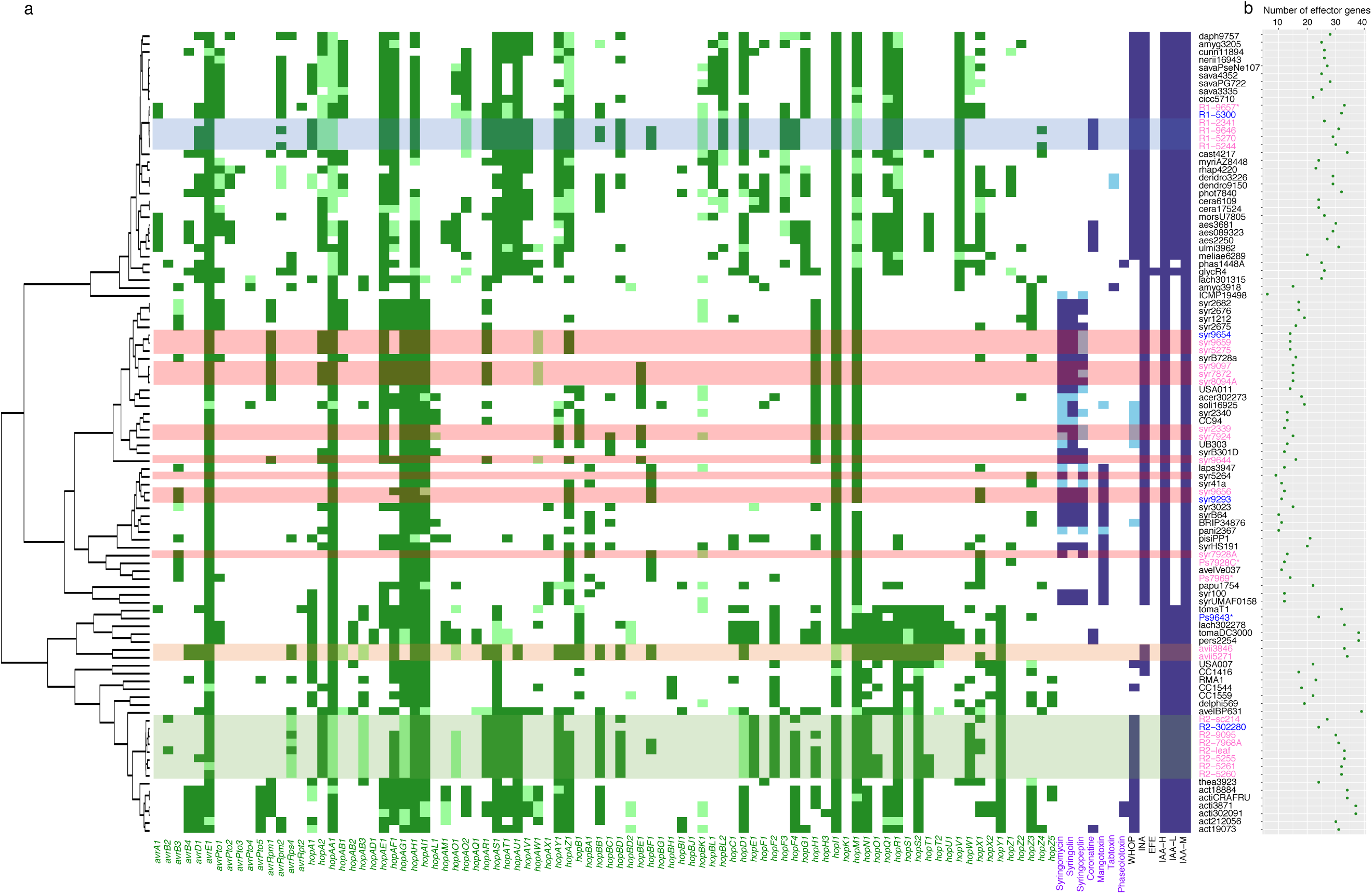
Virulence gene identification. a: Heatmap of virulence gene presence and absence across *P. syringae*. The dark green squares indicate presence of a full-length T3E homologue whereas light green squares indicate that the gene is disrupted or truncated in some way. Other non-T3 secreted virulence factors are coloured in dark and light blue. Strains infecting cherry and plum are highlighted in pink and blue respectively. * indicates non-pathogenic to cherry in controlled pathogenicity tests. The cherry-pathogenic clades are illustrated via horizontal shading of cells with *Psm* R1 in blue, *Psm* R2 in light green, *Pss* in light red and *P.s* pv. *avii* in orange. Strains are ordered based on the core genome phylogenetic tree which is represented by the dendrogram. b: The total number of full-length and pseudogenised T3E genes plotted for each strain.

**Table 3:**
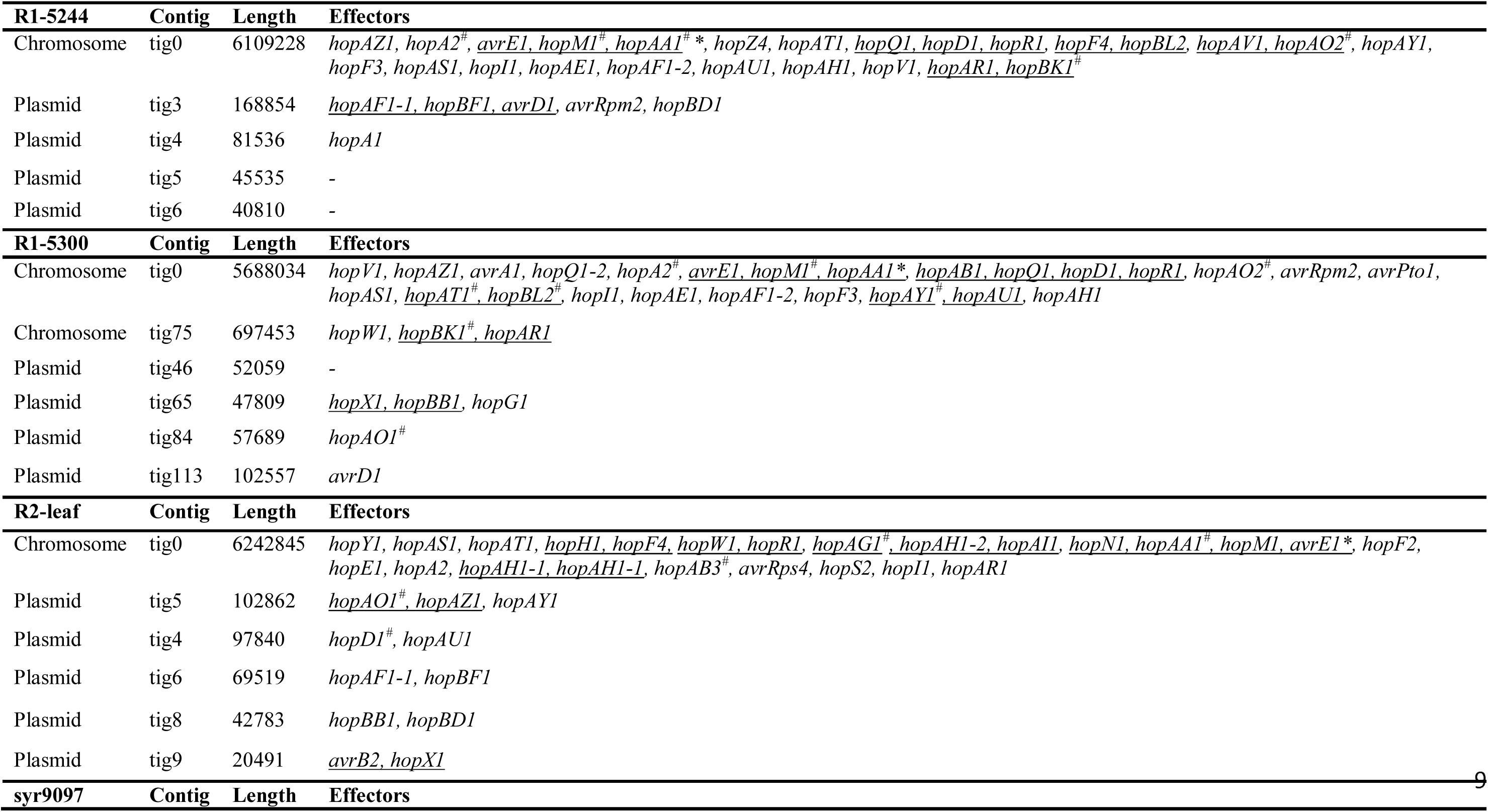
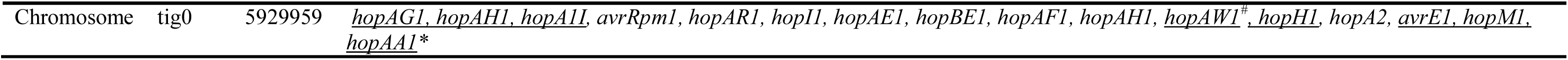
List of effectors in genomes sequenced using PacBio/Minion methods. Effectors are listed in order of appearance on each assembly contig (labelled as chromosomal or plasmid). Where effectors could be considered as linked (within 10kb of each other) they are underlined. * indicates effectors within the conserved effector locus. #: Effector gene is disrupted and is labelled as a pseudogene.

Non-T3 virulence factors were also identified. All pathogenic *Psm* R1 strains possessed the coronatine biosynthesis clusters, which were plasmid-borne in *Psm* R1-5244. All cherry-pathogenic *Pss* strains possessed at least one biosynthesis gene cluster for the toxins syringomycin, syringolin and syringopeptin, with several strains possessing all three. Strains within clade P2b possessed the biosynthesis genes for mangotoxin. The non-pathogenic cherry P2b strains Ps7928C and Ps7969 lacked all toxin biosynthesis clusters.

A cluster of genes named WHOP (<u>w</u>oody <u>h</u>osts and *<u>P</u>seudomonas*) thought to be involved in aromatic compound (lignin) degradation (Caballo-Ponce *et al*., 2016) was present in *Psm* R1 and R2, whilst *P.s* pv. *avii* and most *Pss* strains contained no WHOP homologues. Two cherry P2d strains (syr2339 and syr7924) did however possess the catechol *catBCA* cluster. Finally, the genomes were searched for the ice nucleation gene cluster. Members of *Psm* R1, *Pss* and *P.s* pv. *avii* strains all possessed genes involved in ice nucleation (Fig. 2). Whilst Psm R2 lacked the complete set of genes for ice nucleation.

### Associating T3E evolution with host specificity

T3E evolution was statistically correlated with cherry pathogenicity, using the programs BayesTraits and GLOOME (Pagel, 2004; Cohen *et al*., 2010). BayesTraits takes a binary matrix of two traits and a phylogeny and determines if changes in the two characters (effector gene and pathogenicity) have evolved independently or dependently. Fig. 3a shows the likelihood ratio of cherry pathogenicity being correlated with each effector family’s evolution, with significantly associated effectors highlighted.

**Fig. 3:**
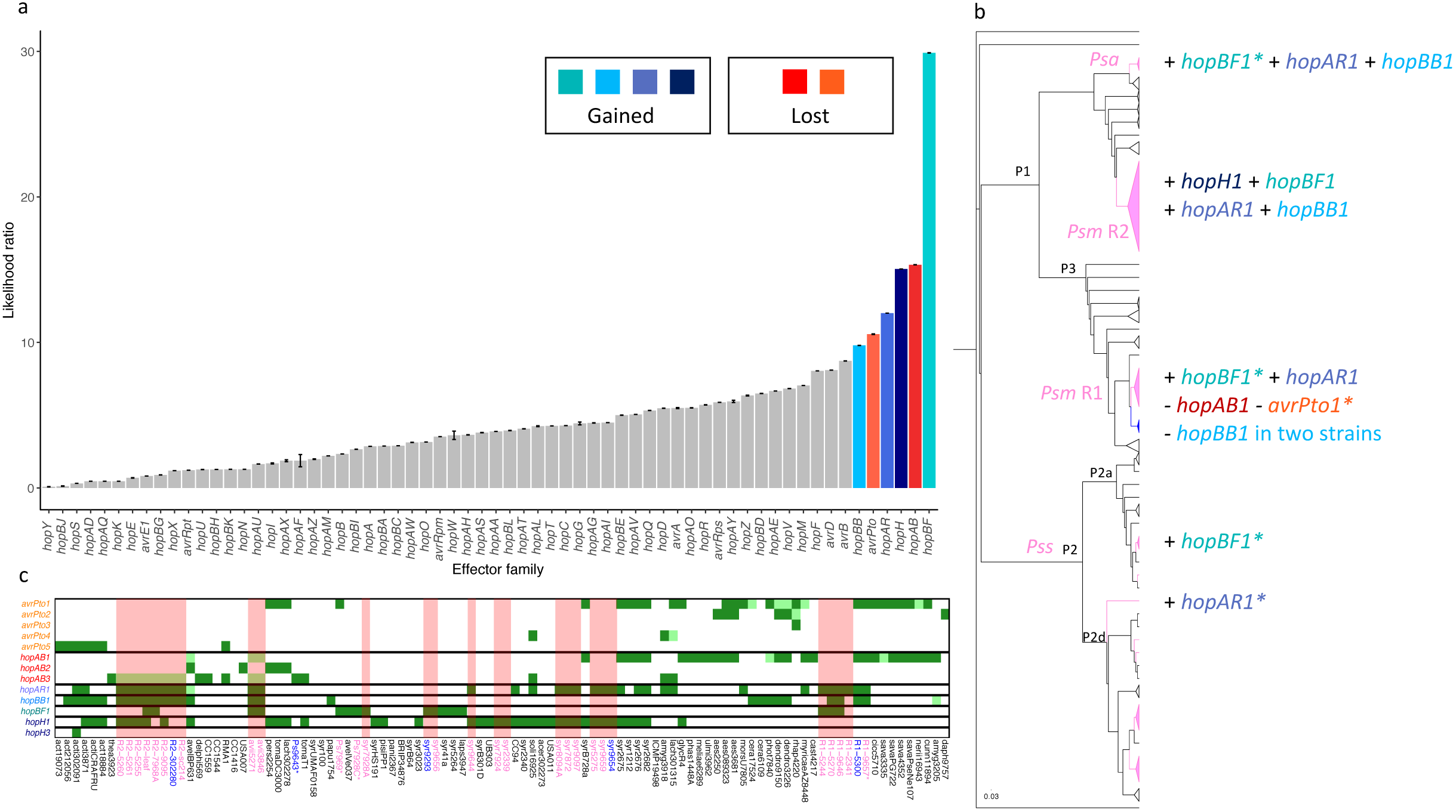
Association of T3E evolution with cherry pathogenicity. a: Barplot showing the likelihood ratio for the correlation of each effector gene family with cherry pathogenicity based on BayesTraits analysis using the core genome phylogeny. The values are obtained from means of 100 independent runs of the program with error bars showing standard error. Those effectors that were not significantly associated with pathogenicity are coloured in grey. Coloured bars were associated with pathogenicity. GLOOME analysis revealed those significant effector genes where presence of the gene was associated with pathogenicity (p≤0.05). These are coloured in shades of blue, whilst where the significant gene was absent in cherry pathogenic clades the bar is coloured in shades of red. b: Gain and loss of BayesTraits-associated T3Es in cherry-pathogenic clades on the core genome phylogeny predicted using GLOOME (probability ≥0.8). For visualisation, clades within the phylogenetic tree have been collapsed with cherry pathogenic clades in pink (*Psm* R1 plum strains in blue). *Psa: P.s* pv. *avii*. Effectors are colour-coded based on the bar colours in A. *: The probability of this effector being gained/lost was slightly lower than 0.8 (see Table S11 for details). c: Heatmap of presence and absence of associated effectors across *P. syringae*. This was constructed as in Fig. 2. Cherry-pathogenic strains are highlighted by pink vertical shading of columns.

BayesTraits analysis using the core genome phylogeny predicted the evolution of six T3E families was linked to cherry pathogenicity. These were *hopBF, hopAB, hopH, hopAR, avrPto* and *hopBB*. To account for any phylogenetic uncertainty, the program was also run on the full set of 100 bootstrapped trees generated by RAxML. The evolution of T3Es *hopBF, hopAR* and *hopAB* was always associated with pathogenicity for all 100 trees, indicating strong association. Whilst, the T3E genes *avrPto, hopBB* and *hopH* were only significantly correlated for 88%, 77% and 62% of trees respectively (Fig. S13). To determine how these genes had been gained or lost across the phylogeny, the program GLOOME was used (Cohen *et al*., 2010). Fig. 3b illustrates the predicted gain and loss of these T3Es on the branches leading to cherry pathogenic clades (with gene presence/absence profiles shown in Fig. 3c). Those putatively associated with pathogenicity (high probability of gain in cherry-pathogenic clades) included *hopAR1, hopBB1, hopBF1* and *hopH1*. The T3Es *hopAB1* and *avrPto1* were found to be lost from cherry pathogenic *Psm* R1, whilst the *hopAB1* and *hopAB3* alleles were pseudogenised in *Psm* R2 and *P.s* pv. *avii* (Fig. 3c). All effector gain and loss events are presented in Fig. S14 and Table S11. Fig. S15 shows the phylogeny with branch labels used in GLOOME.

GLOOME predicted that key effectors have been gained in multiple clades. The *hopAR1* gene has been gained in *Psm* R1, *Psm* R2, *Pss* and *P.s* pv. *avii*. The T3E *hopBB1* was present in the majority of strains within *Psm* R1, R2 and *P.s* pv. *avii* but was absent in *Pss* strains. It showed high probability of gain on branches leading to both *Psm* R2 and *P.s* pv. *avii*. However, GLOOME predicted loss in two *Psm* R1 strains indicating it has experienced dynamic evolution in cherry pathogens. The *hopBB1* effector is closely related to members of the *hopF* family and *avrRpm2* (Lo *et al*., 2016). In addition to the significant acquisition of *hopBB1* homologues, the *hopF* family was expanded in cherry pathogens. Pathogenic strains in *Psm* R1 and R2 all possessed two *hopF* alleles each (*hopF3* and *hopF4/hopF2* and *hopF4*, see Fig. 2). *P.s* pv. *avii* did not possess any *hopF* homologues, but had gained *hopBB1*. By contrast, *Pss* strains lacked all *hopF* members.

### Origins of key effectors in cherry pathogens

To understand the origins of key effectors, gene phylogenies were produced. Incongruence with the core-genome phylogeny indicated that effector sequences had likely experienced HGT between the pathogenic clades, as their sequences clustered together. There has been possible effector exchange between *Psm* R1, R2 and *P.s* pv. *avii*. To predict precisely where transfers had occurred on the phylogeny, the program RANGER-DTL was utilised (Bansal *et al*., 2012). Table 4 reports T3Es that exhibited evidence of HGT between cherry pathogens (gene trees are presented in Fig. S16-S17). Full transfer events are listed in Table S12 and Fig. S18 shows the phylogeny with branch labels used in RANGER-DTL. The BayesTraits correlated T3Es *hopBB* and *hopBF* both showed evidence of HGT. Fig. 4a shows examples of T3Es putatively undergoing HGT between cherry pathogenic clades highlighted in red. Alignments of the flanking regions (Fig. 4b) showed homology between the cherry pathogens and included mobile elements likely involved in recombination events. For example, the regions surrounding *Psm* R1 and R2 *avrD1* sequences were identical (Fig. 4b), indicating that this effector has probably been gained on the same mobile region in both clades. Putatively transferred effectors were mostly plasmid-encoded in the long-read genomes (Table 3). In R1-5244 several of these genes were encoded on one plasmid (Contig 3), whilst in R2-leaf they were found on two plasmids (Contig 6 and 8).

**Table 4:**
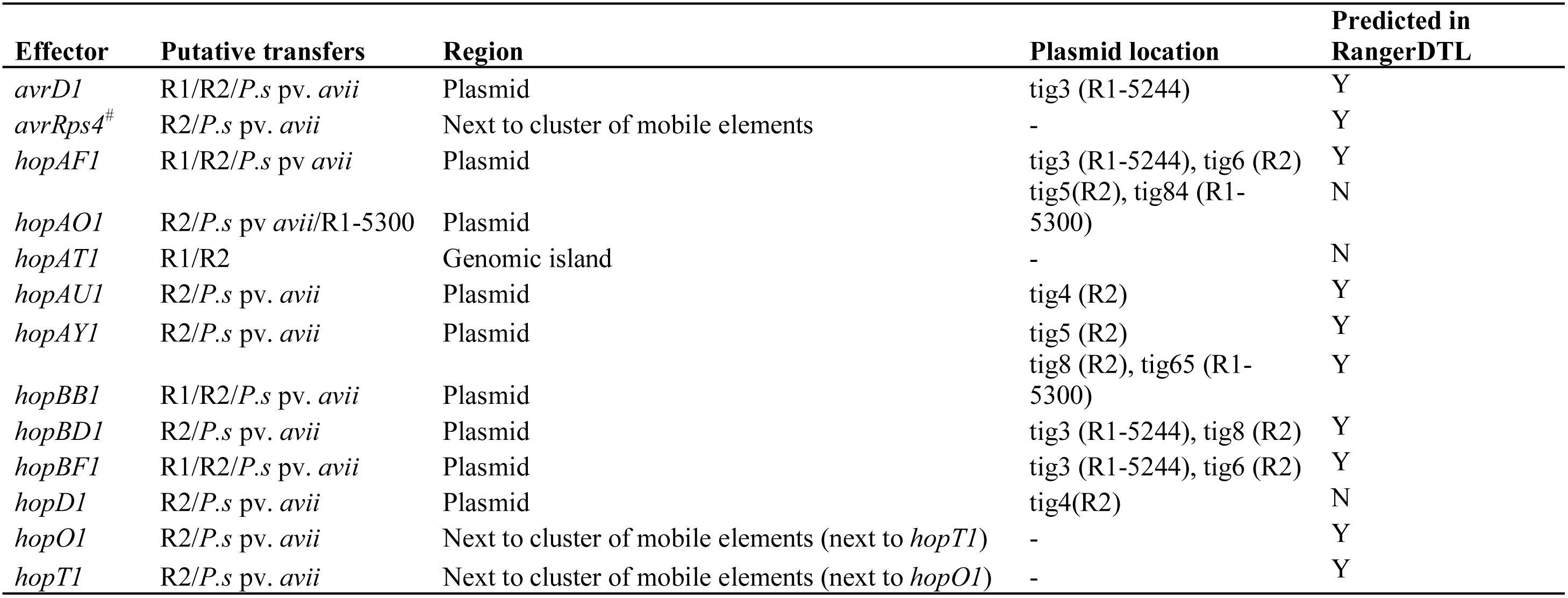
List of putative horizontal gene transfer events that have occurred between Prunus-infecting clades within *P. syringae*. Where the effector gene is present in the PacBio- or Minion-sequenced strains its chromosomal or plasmid location is indicated. #: Effector gene is disrupted and is labelled as a pseudogene.

**Fig. 4:**
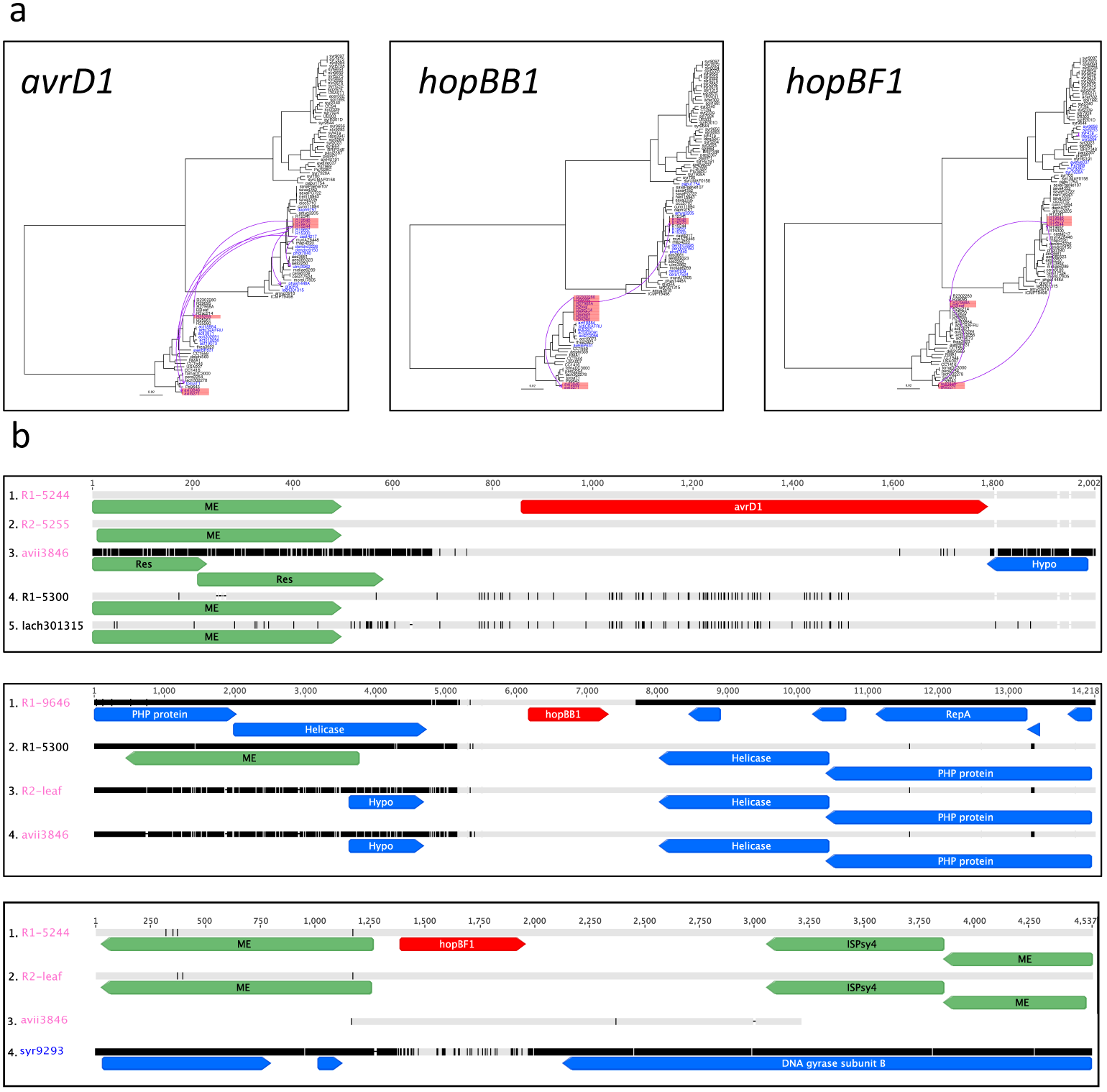
Horizontal gene transfer has played a key role in the evolution of cherry pathogenicity. a: Transfer of *avrD1, hopBB1* and *hopBF1* between different cherry-pathogenic clades based on the *P. syringae* core genome phylogeny. Strains that possess the T3E are coloured in blue, and those that are cherry-pathogenic are highlighted in red. The transfer events predicted by RANGER-DTL are shown by purple arrows. The scale-bar shows substitutions per site. b: DNA alignments of genomic regions containing these effectors. Alignments are colour-coded based on similarity where identical residues are in grey, whereas dissimilar residues appear in black. The effector gene is coloured in red, mobile element genes are in green and other CDS are in blue. Cherry-pathogenic strains are named in pink. Gene name abbreviations: ME, mobile element, Res, resolvase, Hypo, hypothetical protein gene; ISPsy4, insertion sequence, PHP, polymerase and histidinol phosphatase.

The pathogenicity-associated T3E gene *hopAR1* was present in 23/28 cherry pathogens and showed probable gain in pathogenic clades. Phylogenetic analysis of this T3E (Fig. 5a) showed that the sequences for the different cherry pathogenic clades did not cluster with each other, indicating convergent acquisition. Prophage-identification (Table S13) did however reveal that this T3E is predicted to have been gained in *Psm* R1 and R2 within different phage sequences, whilst in *Pss* it is on a genomic island (Fig. 5b), so has been acquired via distinct mechanisms. The *Psm* R1 phage is 51.5kb, described as intact and contains both *hopAR1* and a truncated version of *hopBK1*. The *Psm* R2 phage sequence was 37.1kb and was described as ‘incomplete’, indicating it did not have all the components of an active prophage. Further analysis of this region in *Psm* R2 and P2 strains revealed a shared adjacent tRNA-Thr gene (Fig. 5c,d). Within P2, although cherry *Pss* strains lacked the phage, several strains isolated from bean (syr2675, syr2676 and syr2682) possessed the *hopAR1* gene within a phage homologous to that in *Psm* R2. The syr2675 *hopAR1* sequence was also the most closely related homologue of *Psm* R2 *hopAR1* (Fig. 5a). This novel evidence suggests that this effector gene may have been transferred via phage between phylogroups.

**Fig. 5:**
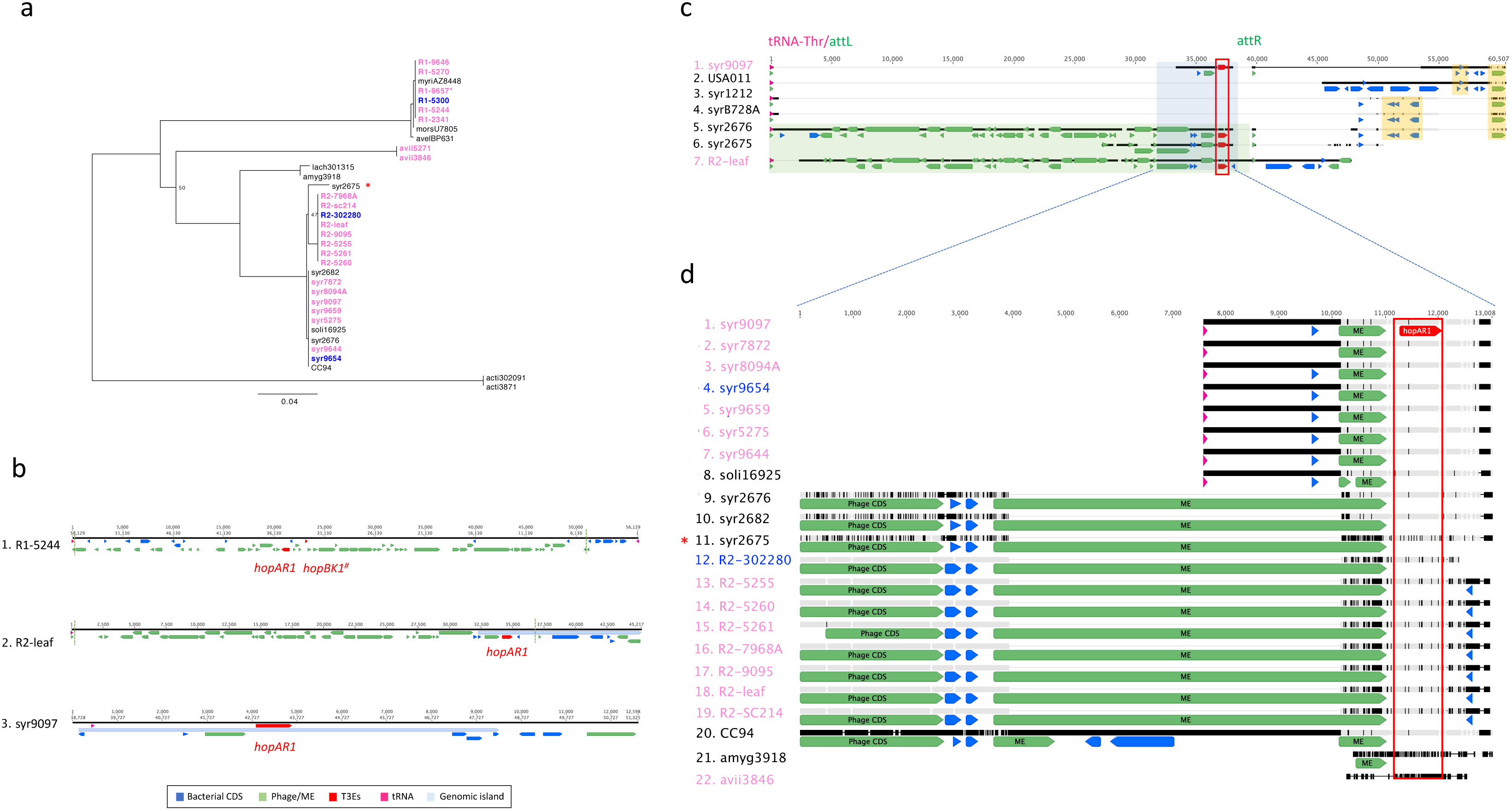
Evolution of *hopAR1* in different clades of *P. syringae* containing cherry pathogens. a: Maximum-likelihood phylogenetic tree built using the nucleotide sequences of the *hopAR1* gene. Cherry and plum isolated strains are highlighted in pink and blue respectively. R1-9657* is classed as a non-pathogen of cherry. Bootstrap supports less than 100% are shown. The scale is nucleotide substitutions per site. * points to the clustering of *Psm* R2 sequences with syr2675. b: Genomic locations of the *hopAR1* gene in the three PacBio-sequenced cherry pathogens. The gene is located within prophage sequences in *Psm* R1 and R2 (see Table S13 for details), whereas in syr9097 it is on a genomic island adjacent to a tRNA gene. Effector genes are coloured in red, other CDS in blue, phage genes predicted by PHASTER and mobile element genes are in green, tRNA genes in pink and genomic islands predicted (GI14 in *Psm* R2 and GI23 in Pss) in light blue. # Indicates that *hopBK1* is a pseudogene in this strain. The end of predicted prophage sequences is denoted with a dashed green line. c: Alignment of the region surrounding the *hopAR1* gene in several strains. The blue box indicates the region in d. All sequences share an upstream tRNA-Thr gene in pink. The phage region in the bean pathogens syr2676 and syr2675 (shortened due to it being at the end of a contig) and *Psm* R2-leaf are highlighted in a green box. Other mobile elements are also coloured in green. The *hopAR1* gene is coloured in red and outlined with a red box. All strains with the phage sequences and syr9097 also contain *attL* and *attR* repeats which are putative insertion sites. Additional closely related phylogroup 2 (P2) strains (USA011, syr1212, syrB728A) which lack the *hopAR1* gene are included for comparison. Homologous regions are highlighted in yellow to show regions of similarity between P2 strains. d: Close-up alignment of the genomic regions surrounding the *hopAR1* gene in strains of P2, *Psm* R2, *P.s* pv. *amygdali* and *P.s* pv. *avii* that share homology in the surrounding regions. tRNA genes are in pink, phage/mobile element genes are in green, CDS in blue and the *hopAR1* T3E gene is coloured red and highlighted with a red box. Alignments are colour-coded based on similarity where identical residues are in grey, whereas dissimilar residues appear in black. * The syr2675 *hopAR1* gene is similar to *Psm* R2 sequences and also has homologous phage sequences upstream.

Many T3Es are mobilised between bacteria on genomic islands (GI). GIs were identified for the three PacBio-sequenced strains of *Psm* R1, *Psm* R2 and *Pss* (Tables S14-S16). R1-5244 GIs contained the coronatine biosynthesis cluster and six T3Es. In R2-leaf, eight T3E genes were located on GIs, whilst in syr9097 three T3Es were found on genomic islands.

These GIs were then searched for in other *P. syringae* genomes to identify potential sources of transfer and Fig. 6a shows heatmaps of GI presence. The *Psm* R1 GIs included several found only in pathogenic *Psm* R1 strains differentiating them from the non-pathogens. These included the coronatine biosynthesis cluster (GI1), *hopF3* (GI6) and *hopAT1* (GI14). Most *Psm* R1 GIs produced hits across *P. syringae*, particularly in P1 and P3. *Psm* R2 GIs were most commonly found in P1. Several were shared with other cherry-pathogenic clades, including those containing *hopAF1* (GI36), *hopAT1* (GI3) and *hopD1* (GI6). Finally, although most islands identified in syr9097 were commonly found across the species complex, those containing T3Es (GI30, GI23 and GI26) appeared to be P2-specific, indicating that cherry-pathogenic strains likely gained these islands from other members of P2.

**Fig. 6:**
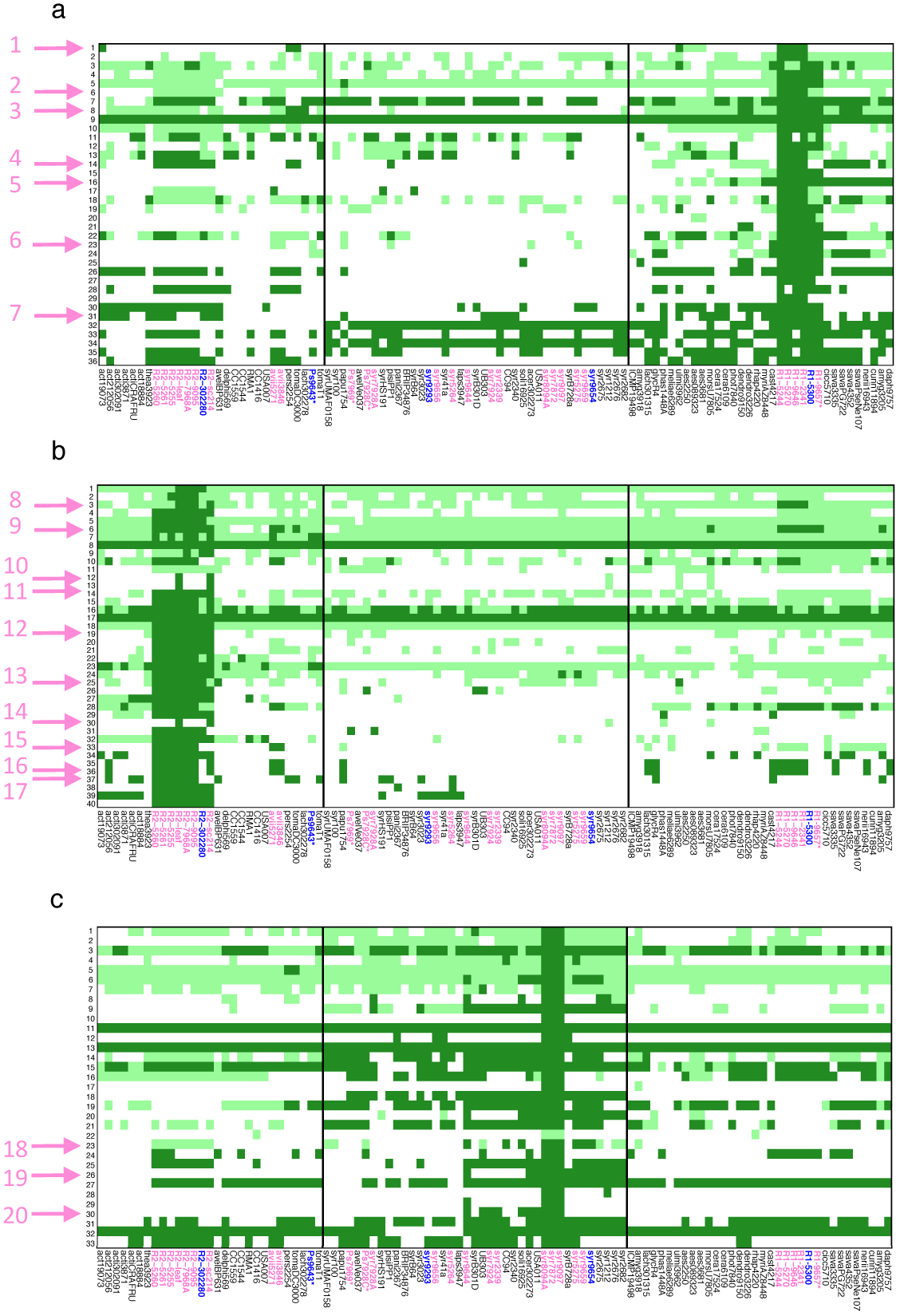
Genomic islands characteristic of cherry pathogens are found across *P. syringae*. The heatmap shows the presence and absence of the genomic islands identified in the PacBio sequenced pathogenic strains of (a) *Psm* R1, (b) *Psm* R2 and (c) *Pss* across the *P. syringae* complex. Dark green squares indicate that the full-length GI is putatively present. Light green squares are where the GI was predicted to be partially present. The names of strains that infect cherry and plum are coloured in pink and blue, respectively. Strains that were not pathogenic but were isolated from cherry are marked with an asterisk. The different phylogroups 1,2 and 3 are delimited by black borders. Pink arrows point to GIs that contain T3E or phytotoxin genes. PsmR1 GIs – 1: coronatine biosynthesis genes, 2: *hopF3*, 3: *hopA1*, 4: *hopAT1*, 5: *hopBL2*, 6: *hopAO2*, 7: *hopAY1, Psm* R2 GIs - 8: *hopAT1*, 9: *hopD1*, 10: *hopX1*, 11: *hopAR1*, 12: *hopE1*, 13: *hopAE1*, 14: *avrB2*, 15: *hopAZ1*, 16: *hopAF1*, 17: *hopH1, Pss* GIs - 18: *hopAR1*, 19: *avrRpm1*, 20: *hopBE1*. Full details are in Tables S14-S16.

### Functional analysis of potential *avr* genes

To validate the predictive power of this analysis, cloning was used to identify avirulence factors active in cherry. The effector genes *avrPto* and *hopAB* were absent from cherry pathogens and their evolution was theoretically linked to pathogenicity. Several other candidate avirulence effectors were identified, that were absent from cherry pathogens, but present in close out-groups (Fig. 2). Avirulence-gene identification focused on *Psm* R1 as any T3E variation within the clade may be due to differences in host specificity rather than phylogenetic distance. Potential avirulence T3E genes included *avrA1, avrPto1, hopAA1, hopAB1, hopAO2* and *hopG1*, which had full-length homologues in non-pathogenic *Psm* R1 strains, but were absent from or truncated in pathogens. These genes were cloned from R1-5300 (except *hopAO2*, which was cloned from R1-9657).

The effector *avrRps4* was also cloned from *P.s* pv. *avellanae* (*Psv*) BPIC631 a close relative of *Psm* R2. This effector was absent from most cherry-pathogenic strains. Several pathogens possessed the full-length gene (R2-leaf, R2-9095 and *P.s* pv. avii), but lacked the KRVY domain that functions *in planta* (Fig. S19) (Sohn *et al*., 2009). The *hopAW1* gene was cloned from *Pph*1448A as this T3E has undergone two independent mutations in *Pss* strains, disrupting the beginning of the gene (Fig. S20). Finally, *hopC1* was cloned from the *Aquilegia vulgaris* pathogen RMA1 which is basal to the *Psm* R2 clade, as it is absent from all cherry-pathogenic strains.

Nine effectors were cloned into the expression vector pBBR1MCS-5 and conjugated into three pathogenic strains (R1-5244, R2-leaf and syr9644). Knock-out strains for the T3SS gene *hrpA* were obtained for R1-5244 and R2-leaf to act as non-pathogenic controls that could not secrete T3Es and failed to cause the HR on tobacco (Fig. S21).

Bacterial population counts were conducted in cherry leaves. The transconjugants expressing HopAB1 or HopC1 failed to multiply to the same levels as the pathogenic empty vector (EV) controls or produce disease lesions. The expression of AvrA1, AvrRps4, and HopAW1 also caused significant reductions in population growth, but this reduction was not consistently seen across all three pathogenic strains (Fig. 7a). As the addition of the *hopAB1* gene reduced pathogenicity, the *hopAB2* and *hopAB3* genes were also cloned from PsvBPIC631 and RMA1, and were also found to reduce pathogen multiplication (Fig. 7b).

**Fig. 7:**
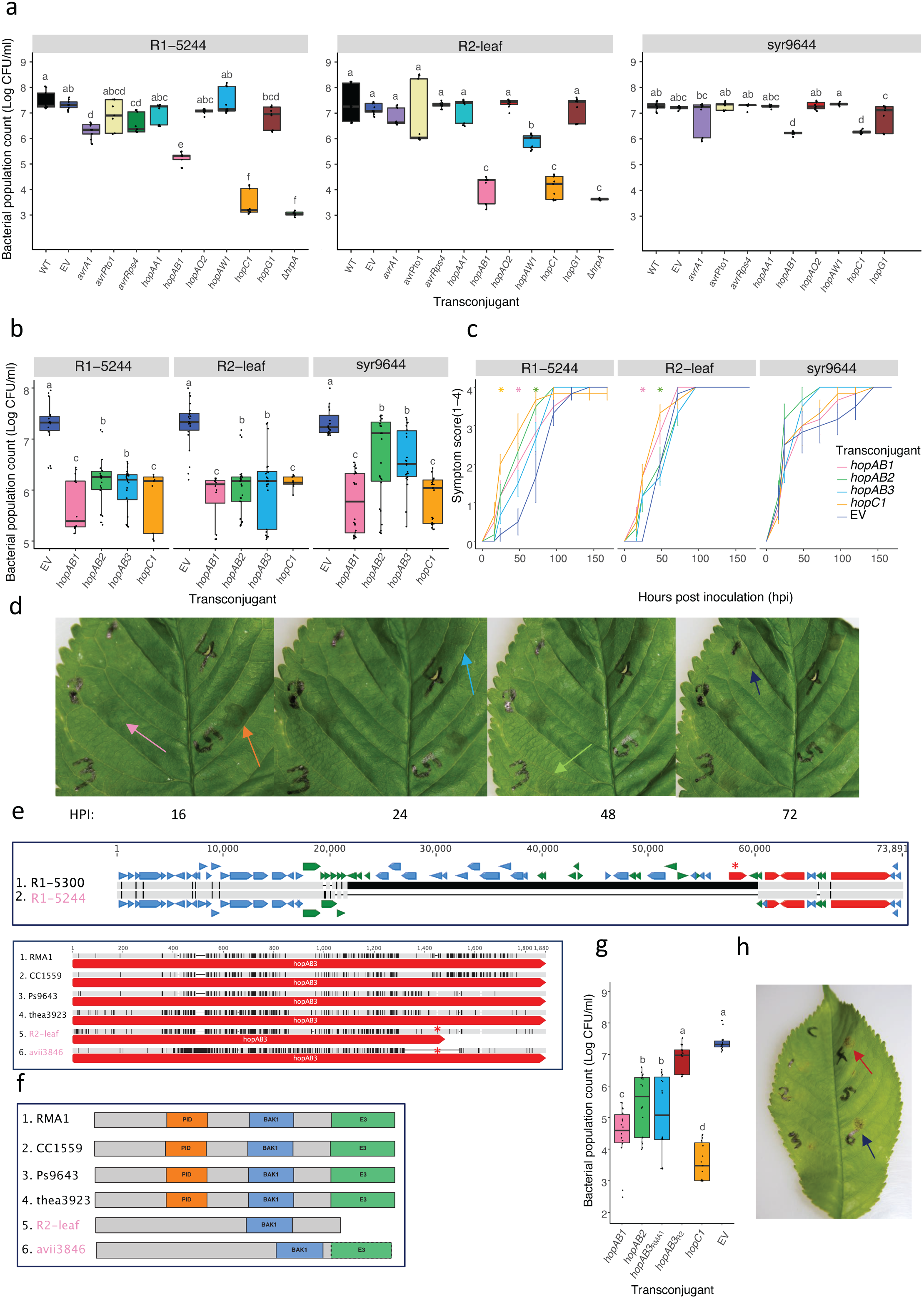
Identification of avirulence factors activating ETI in cherry. a: Boxplot of an initial ten-day population count analysis of cherry pathogens (R1-5244, R2-leaf and syr9644) transconjugants expressing candidate avirulence genes. The data presented are based on one experiment, with three leaf replicates and three nested technical replicates (n=9). Controls included the wildtype strain, a strain containing the empty pBBR1MCS-5 vector and a *ΔhrpA* deletion mutant (for R1-5244 and R2-leaf). A separate ANOVA was performed for each cherry pathogen (R1-5244, R2-leaf and syr9644) and the Tukey-HSD significance groups (p=0.05, confidence level: 0.95) for each strain are presented above each boxplot. b: Boxplot of ten-day population counts of cherry pathogens (R1-5244, R2-leaf and syr9644) expressing different HopAB alleles and HopC1. The data presented are based on three independent experiments (n=27). Tukey-HSD significance groups (p=0.05, confidence level: 0.95) are presented above each boxplot. c: Symptom development of R1-5244, R2-leaf, syr9644 transconjugants. Mean symptom score values are presented and represent two independent experiments (n=6). Error bars show standard error above and below the mean. Symptoms assessed as degree of browning of the infiltration site, 1: some browning, 2:< 50%, 3:>50%, 4: 100% of the infiltrated area brown. Analysis was based on Area Under Disease Progress Curves values (0-48 h), see Tables S25-S26. An ANOVA was performed on AUDPC values, with * indicating significantly different from the empty vector (EV) control. d: Symptom development over time on a leaf inoculated with R1-5244 transconjugants. HPI: Hours post inoculation. The order of strains: 1: EV, 2: *hopAB1*, 3: *hopAB2*, 4: *hopAB3*, 5: *hopC1*. Arrows indicate the first appearance of symptoms associated with each strain and are coloured based on the graph in c. e: Alignment of the DNA region surrounding *hopAB1* in *Psm* R1 strains. Grey indicates sequence identity whereas black indicates divergence. The effector genes are coloured in red, whereas other CDS are in blue and putative mobile element genes are in green. * indicates the location of *hopAB1* in R1-5300, whilst the upstream effectors are *hopQ1, hopD1* and *hopR1*. The second alignment shows the *hopAB3* gene of *Psm* R2 and close out-groups. * The *hopAB3* gene has been truncated due to a GG insertion leading to a frameshift in *Psm* R2, whilst in *P.s* pv. *avii* (avii3846) there is a deletion at the end of the gene. f: Diagrams showing the location of key domains in the HopAB3 protein including the Pto-interaction domain (PID), BAK1-interacting domain (BAK1) and E3 ubiquitin ligase (E3). The E3 domain is lost completely from the *Psm* R2 protein whereas in avii3846 the beginning of this domain is lost. The PID domain was not detected in the cherry pathogen sequences. g: Boxplot of ten-day population counts of R1-5244 trans-conjugants expressing three different *hopAB* alleles, including *hopAB3_R2-leaf_* and *hopC1*. The data presented are based on two independent experiments (n=18). Tukey-HSD significance groups (p=0.05, confidence level: 0.95) are presented above each boxplot. h: Representative image of symptoms 10 dpi with the different R1-5244 transconjugants when inoculated at a low level to observe pathogenicity. Arrows point to pathogenic symptoms in the strain expressing *hopAB3_R2-leaf_* and the EV strain, colour-coded as in g. ANOVA tables for all statistical analyses are presented in Tables S17-S25.

To further investigate the induction of the HR by the HopAB family and HopC1, inoculations were performed at high concentrations (2x10^8^ CFU/ml) as described in Hulin *et al*. (2018). In *Psm* R1 and R2, the addition of these T3Es led to more rapid tissue collapse than observed in EV controls, indicative of HR induction (Fig. 7c,d); HopC1 and HopAB1 were particularly effective. With *Pss*, however, EV transconjugants themselves caused rapid tissue collapse, making it impossible to recognise an induced HR as symptom development was not significantly different.

The *hopAB1* gene is found in a mobile-element rich ~40kb region in the non-pathogenic *Psm* R1-5300, missing from the pathogen *Psm* R1-5244 (Fig. 7d). Meanwhile, *Psm* R2 and *P.s* pv. *avii* possessed putatively pseudogenised *hopAB3* alleles (Fig. 7e) and *P.s* pv. *avii* possessed a truncated *hopAB1* gene (Fig. S22). *hopAB3* is truncated in *Psm* R2 due to a 2bp insertion (GG at position 1404bp) leading to a premature stop codon, whilst in *P.s* pv. *avii* a 218bp deletion has disrupted the C-terminus. If expressed, the E3-ubiquitin ligase is completely absent from the *Psm* R2 protein and disrupted in *P.s* pv. *avii* (Fig. 7f). Both HopAB3 alleles were also divergent enough that the Pto-interacting domain (PID) was not identified by Interproscan. To determine if the truncated *Psm* R2 HopAB3 allele induced any resistance response in cherry leaves, the gene was expressed in *Psm* R1-5244 and population growth measured. The addition of this gene did not lead to a significant reduction in growth compared to the EV control, unlike other *hopAB* alleles (Fig. 7g) and the transconjugant was still able to induce disease symptoms 10 dpi (Fig. 7h).

## Discussion

### Core genome phylogenetics

Phylogenetic analysis confirmed that cherry pathogenicity has evolved multiple times within *P. syringae. Psm* R1, R2 and *P.s* pv. *avii* each formed distinct monophyletic clades, whereas cherry-pathogenic *Pss* strains were distributed across the P2 clade thus indicating that cherry pathogenicity has either evolved multiple times within P2 or that this clade is not highly specialised. To confirm this genomic prediction of pathogenicity, several additional P2 strains isolated from bean, pea and lilac were tested for pathogenicity in cherry. They each produced lower population levels in cherry leaves than cherry pathogens, suggesting that host adaptation has occurred (Fig. S10). Many P2 strains have previously been named *Pss* on the basis of lilac pathogenicity, despite being pathogenic to other plant species (Young, 1991). A new naming system within this phylogroup would therefore be desirable.

### Search for candidate effectors involved in cherry pathogenicity

Gains and losses of T3Es were closely associated with pathogenicity. Virulence-associated effectors *hopAR1, hopBB1, hopH1* and *hopBF1* had been gained in multiple cherry-pathogenic clades. The *hopAR1* effector has been studied in the bean pathogen *P.s* pv. *phaseolicola* R3 (1302A), as an GI-located *avr* gene (*avrPphB*) whose protein is detected by the corresponding R3 resistance protein *in planta* (Pitman *et al*., 2005; Neale *et al*., 2016). HopAR1 also acts as a virulence factor as a cysteine protease which targets receptor-like kinases to interfere with plant PAMP-triggered immunity (PTI) responses (Zhang *et al*., 2010). This effector could play a similar role in PTI suppression in cherry.

HopBB1 and other members the HopF family were abundant in cherry pathogens. All HopF members share an N-terminus and myristoylation sites for plant cell membrane localisation (Lo *et al*., 2016) and interfere with PTI and Effector-triggered immunity (ETI) in model plants (Wang *et al*., 2010; Wu *et al*., 2011; Hurley *et al*. 2014). The presence of multiple *hopF* homologues in cherry pathogens and specific gain of *hopBB1* suggested the importance of their function. In comparison, HopH1 and HopBF1 are under studied. HopH1 is a protease, homologous to the *Ralstonia solanacearum* Rip36 protein (Nahar *et al*., 2014). This T3E gene was found on GI37 in *Psm* R2-leaf and was within 3kb of *hopF4* (Fig. S23), indicating that these two T3Es may have been gained together. HopBF1 was first discovered in *P.s* pvs *aptata* and *oryzae* (Baltrus *et al*., 2011) but its role *in planta* is undetermined. This study therefore provided candidate T3Es important for cherry pathogenicity which could be the focus of future functional studies.

Phytotoxin biosynthesis gene clusters were also identified. Coronatine has been gained on a plasmid in pathogenic *Psm* R1 and could be one of the factors that differentiate pathogens from non-pathogens in this clade. Coronatine functions in virulence by down-regulating salicylic acid defence signalling (Grant & Jones, 2009). Necrosis-inducing lipodepsipeptide toxins were common in P2. All cherry-pathogenic *Pss* strains possessed at least one biosynthesis cluster. The ability of *Pss* strains to cause necrosis on cherry fruits has been linked to toxins (Scholz-Schroeder *et al*., 2001). Interestingly, two non-pathogenic P2b cherry strains lacked all phytotoxins, a deficiency that could contribute to their lack of pathogenicity.

All cherry-pathogenic *Pss* strains had reduced effector repertoires. This observation supports the hypothesis that a phenotypic trade-off exists, with strains retaining few T3Es, whilst relying more on phytotoxins for pathogenicity (Hockett *et al*., 2014). If this pathogenic strategy has evolved in the P2 clade, this raises the question as to how it affects host specificity and virulence. P2 strains often infect more than one host species (Rezaei & Taghavi, 2014). These strains probably possess fewer ETI-inducing avirulence factors that restrict effector-rich strains to particular hosts, so may be more successful generalists. The reduction in T3E repertoire could however be limiting as strains may be less capable of long-term disease suppression required at the start of a hemi-biotrophic interaction.

Most cherry-pathogenic clades possessed genes involved in aromatic compound degradation, shown to be important in virulence on olive (Caballo-Ponce *et al*., 2016) and ice nucleation genes that stimulate frost damage (Lamichhane *et al*., 2014). The fact that not all cherry-pathogenic clades possessed these genes suggests they are not crucial for bacterial canker, however they could contribute to niche persistence. For example, Crosse and Garrett (1966), observed that *Psm* R1 survived in cankers for longer than *Pss*. Increased persistence could be linked to genes involved in woody-tissue adaptation.

### HGT has been important in the acquisition of key effectors

HGT is a key mechanism for effector shuffling within *P. syringae* (Arnold & Jackson, 2011). Pathogenicity-associated T3Es *hopBB1* and *hopBF1* were plasmid encoded and showed evidence of HGT between the cherry-pathogenic clades in P1 and P3. Plasmid profiling revealed that cherry pathogens in phylogroups 1 and 3 possessed native plasmids, some of which were putatively conjugative, indicating the importance of plasmids in gene exchange. By contrast, most cherry-pathogenic *Pss* strains lacked plasmids.

The pathogenicity-associated T3E *hopAR1* was chromosomal in *Psm* R1 and R2. This gene was found within distinct prophage sequences in these two clades. To our knowledge this is the first reported example of a plant pathogen T3E located within a prophage sequence. Interestingly, the *Psm* R2 *hopAR1* gene homologue was most similar to *hopAR1* from a P2 bean strain syr2675, which is a close relative of cherry *Pss*. This strain possessed the same phage as *Psm* R2, indicating that horizontal gene transfer of this T3E between phylogroups may have been phage-mediated. The striking example of convergent acquisition of *hopAR1* in the cherry pathogens, putatively through distinct prophages in *Psm* R1 and R2, and a GI in *Pss* indicates that this T3E may have important roles in virulence. The well characterised *P.s* pv. *phaseolicola* R3 homologue is not associated with a phage, but has been shown to undergo dynamic evolution on a mobile genomic island *in planta* in resistant bean cultivars (Neale *et al*., 2016). It is intriguing that the same T3E may be important in a completely different pathosystem.

Several T3Es in *Psm* R1, R2 and *Pss* were located on GIs. To determine the likely source of GIs in cherry strains, all other *P. syringae* strains were searched for homologous sequences (Fig. 6). There was evidence of *Psm* R1 and R2 islands being shared between cherry pathogen clades indicative of HGT events occurring between strains occupying the same ecological niche.

### Functional genomics revealed convergent loss of an avr factor

Genes from the *hopAB* and *avrPto* families form a redundant effector group (REG) vital for early PTI suppression in herbaceous species (Jackson *et al*., 1999; Lin & Martin, 2005; Kvitko *et al*., 2009). Both effectors also trigger ETI by interacting with the serine-threonine kinase R protein Pto in tomato (Kim *et al*., 2002).

Across the *P. syringae* complex, the REG was common (Fig. S24), but cherry pathogens all lacked full-length members of this family. The *hopAB1* gene has been lost from *Psm* R1, whilst the *Psm* R2 and *P.s* pv. *avii* predicted HopAB3 proteins lacked the PID and E3-ubiquitin ligase domains through contrasting mutations. *P.s* pv. *avii* also possessed a truncated *hopAB1* gene (Fig. S22), lacking the PID domain. The lack of a PID in the cherry pathogen HopAB proteins suggested that they could have diverged to avoid a Pto-like recognition in cherry.

Full-length members of this REG were cloned into cherry pathogens to determine their role *in planta*. The addition of HopAB alleles (HopAB1-3) consistently reduced population growth of pathogenic strains *in planta* and triggered an HR. HopAB loss or pseudogenisation in cherry pathogens may have been selected for to reduce avirulence activity. The truncated version of HopAB3 in R2-leaf was found not to exhibit avirulence activity as its expression did not reduce the growth of R1-5244 *in planta*. Although AvrPto is part of the same REG, its expression had no effect on the ability of cherry pathogens to multiply *in planta*. The absence of AvrPto in cherry pathogens is therefore unlikely to be driven by avirulence, but could be due to the lack of HopAB virulence targets *in planta*. As this REG is vital for early disease suppression in model strains, cherry pathogens may rely on other T3Es to fulfil this role.

The variation in *hopAB1* presence in *Psm* R1 is intriguing. *Psm* R1 strains may be pathogenic on both cherry and plum (Δ*hopAB1*) or just pathogenic on plum (possessed *hopAB1*) (Hulin *et al*., 2018). This suggested that the host proteins in cherry that detect the presence of HopAB are not present/functioning in plum. Future studies may determine how the two host immune responses diverged and could examine *hopAB* diversity across *Prunus* pathogens. This study focused on *Prunus avium*, however strains isolated from additional *Prunus* spp. were included, such as *P.s* pv. *cerasicola, P.s* pv. *morsprunorum* FTRSU7805, *P.s* pv. *amygdali* and *P.s* pv. *persicae* (Table 1). All apart from *P.s* pv. *amygdali* 3205 and *P.s* pv. *persicae* lacked HopAB (Fig. 2), indicating that there may be a conserved resistance mechanism regulating ETI activated by this effector family in *Prunus* species.

### Linking genomics to host specialisation

Cherry pathogenicity has arisen independently within *P. syringae*, with strains using both shared and distinctive virulence strategies. Cherry-pathogenic clades in P1 and P3 have large effector repertoires. Cherry *Pss* were found across P2. They possessed reduced T3Es and several phytotoxin gene clusters. Key events in the evolution of cherry pathogenicity (Fig. 8) appear to be the acquisition of virulence-associated effectors, often through horizontal gene transfer, such as *hopAR1*, members of the *hopF* family such as *hopBB1, hopBF1* and *hopH1*. Significantly, the loss/pseudogenisation of HopAB effectors has also occurred in multiple clades. Within P2, the different cherry-infecting *Pss* clades have slight differences in their virulence factor repertoires which may reflect their convergent host adaptation. Clades differed in T3E content, phytotoxin genes and possession of genes for catechol degradation. This study demonstrated that populations genomics can be used to examine a complex disease of a perennial plant species. A huge dataset was narrowed down to several candidate host-specificity controlling genes, two of which (*hopAB* and *hopC1*) encode proteins that had putative avirulence functions *in planta*.

**Fig. 8:**
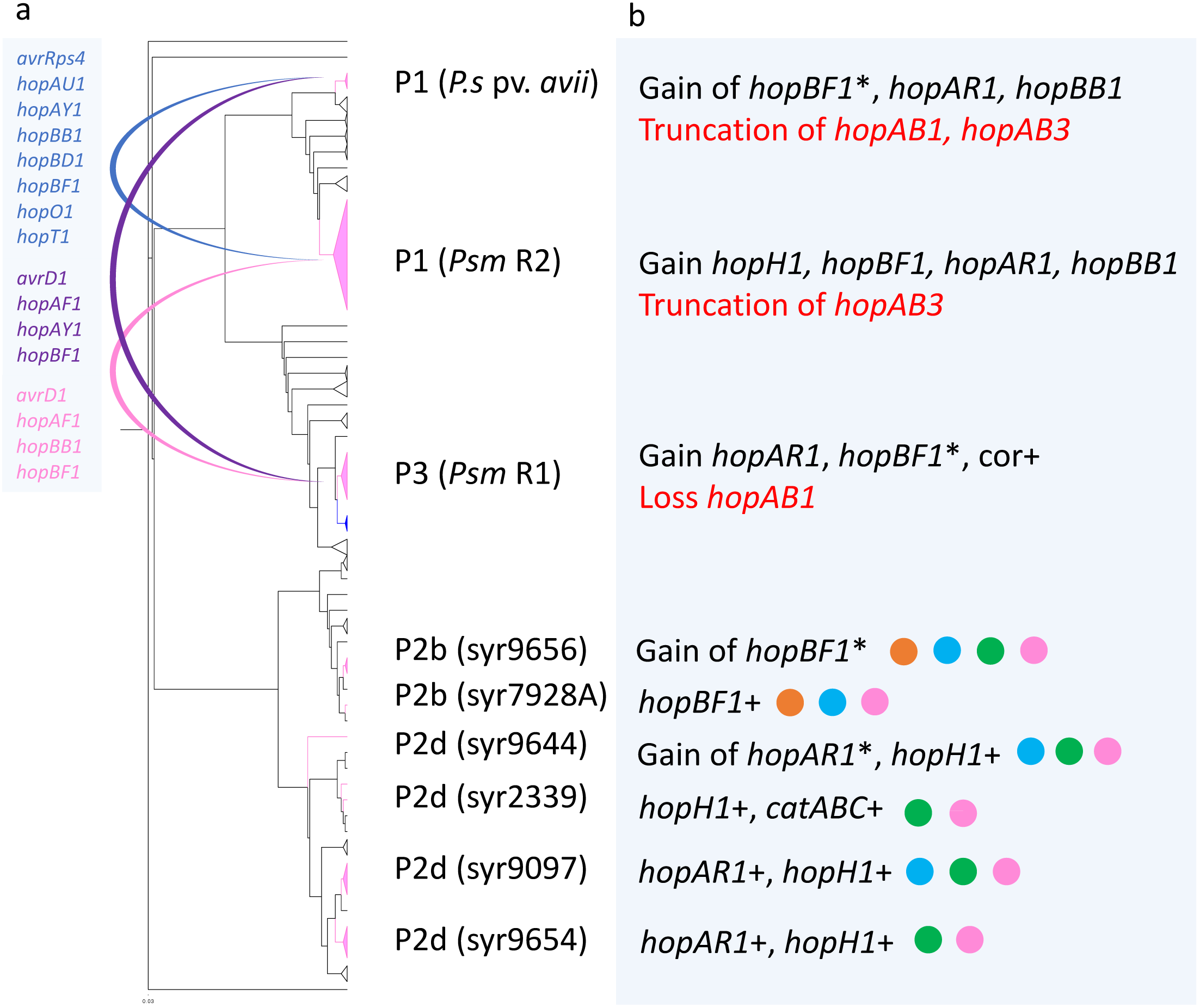
Model highlighting genomic events that have led to the evolution of pathogenicity towards cherry. a: The core genome phylogeny is presented. Scale bar shows substitutions per site. For visualisation, clades within the phylogenetic tree have been collapsed with clades containing cherry pathogens in pink (*Psm* R1 plum strains in blue). Examples of cherry pathogens within each clade of phylogroup 2 are named. HGT events predicted using phylogenetic analysis and RANGER-DTL are shown. b: The key gains and losses of associated virulence genes in strains pathogenic to cherry are described. Where a gene is present but not necessarily predicted to be gained in this clade, it is shown with a plus sign, whilst the use of the words gain or loss specifically denotes results based on GLOOME analysis. *: The probability of this effector being gained/lost predicted using GLOOME was slightly lower than 0.8 (see Table S11 for details). For *Pss* strains within P2, present toxin biosynthesis gene clusters are shown as dots for comparison. Orange: mangotoxin, blue: syringopeptin, green: syringolin, pink: syringomycin.

## Acknowledgements

We acknowledge the East Malling Trust, University of Reading and BBSRC for funding (BB/P006272/1). Library preparation for PacBio sequencing was performed at the Earlham Institute, whilst for wild cherry strains Illumina MiSeq sequencing, libraries was prepared at the Genomics Facility, University of Warwick. We thank Steve Roberts, Helen Neale, Mateo San José and David Guttman for providing bacterial strains. We thank the East Malling Farm and Glass staff for plant maintenance. The authors declare no conflict of interest.

## Author contributions

MTH, JWM, RWJ and RJH conceived and designed the study as well as writing the manuscript. MTH performed bioinformatics, statistical analysis and laboratory work. JV isolated some of the strains used in this study, JV and EH selected representative strains and prepared DNA for MiSeq sequencing of nine strains and LB assembled these nine sequences. HJB performed the MinION library preparation and sequencing. ADA assisted in bioinformatics pipeline development. All authors read and reviewed the final manuscript.

## References

Alfano JR, Charkowski A, Deng W, Badel J, Petnicki-Ocwieja T, van Dijk K, Collmer A. 2000. The *Pseudomonas syringae* Hrp pathogenicity island has a tripartite mosaic structure composed of a cluster of type III secretion genes bounded by exchangeable effector and conserved effector loci that contribute to parasitic fitness and pathogenicity in pl. Proceedings of the National Academy of Sciences of the United States of America 97: 4856–4861.

Almeida NF, Yan S, Lindeberg M, Studholme DJ, Schneider DJ, Condon B, Liu H, Viana CJ, Warren A, Evans C, et al. 2009. A draft genome sequence of *Pseudomonas syringae* pv. *tomato* T1 reveals a type III effector repertoire significantly divergent from that of *Pseudomonas syringae* pv. *tomato* DC3000. Molecular Plant-Microbe Interactions: MPMI 22: 52–62.

Altschul SF, Gish W, Miller W, Myers EW, Lipman DJ. 1990. Basic local alignment search tool. Journal of molecular biology 215: 403–410.

Arndt D, Grant JR, Marcu A, Sajed T, Pon A, Liang Y, Wishart DS. 2016. PHASTER: A better, faster version of the PHAST phage search tool. Nucleic acids research 44: W16–W21.

Arnold DL, Gibbon MJ, Jackson RW, Wood JR, Brown J, Mansfield JW, Taylor JD, Vivian A. 2001. Molecular characterization of *avrPphD*, a widely-distributed gene from *Pseudomonas syringae* pv. *phaseolicola* involved in non-host recognition by pea (*Pisum sativum*). Physiological and Molecular Plant Pathology 58: 55–62.

Arnold DL, Jackson RW. 2011. Bacterial genomes: Evolution of pathogenicity. Current opinion in plant biology 14: 385–391.

Aziz RK, Bartels D, Best AA, DeJongh M, Disz T, Edwards RA, Formsma K, Gerdes S, Glass EM, Kubal M, et al. 2008. The RAST Server: Rapid annotations using subsystems technology. BMC genomics 9: 75.

Baltrus DA, Nishimura MT, Romanchuk A, Chang JH, Mukhtar MS, Cherkis K, Roach J, Grant SR, Jones CD, Dangl JL. 2011. Dynamic evolution of pathogenicity revealed by sequencing and comparative genomics of 19 *Pseudomonas syringae* isolates. PLoS Pathogens 7: 22.

Baltrus DA, Nishimura MT, Dougherty KM, Biswas S, Mukhtar MS, Vicente JG, Holub EB, Dangl JL. 2012. The molecular basis of host specialization in bean pathovars of *Pseudomonas syringae*. Molecular Plant-Microbe Interactions MPMI 25: 877–888.

Baltrus DA, Yourstone S, Lind A, Guilbaud C, Sands DC, Jones CD, Morris CE, Dangl JL. 2014a. Draft genome sequences of a phylogenetically diverse suite of *Pseudomonas syringae* strains from multiple source populations. Genome Announcements 2: e01195–13.

Baltrus DA, Dougherty K, Beckstrom-Sternberg SM, Beckstrom-Sternberg JS, Foster JT. 2014b. Incongruence between multi-locus sequence analysis (MLSA) and whole-genome-based phylogenies: *Pseudomonas syringae* pathovar *pisi* as a cautionary tale. Molecular plant pathology 15: 461–465.

Baltrus DA, McCann HC, Guttman DS. 2017. Evolution, genomics and epidemiology of *Pseudomonas syringae*. Molecular Plant Pathology 18: 152–168.

Bansal MS, Alm EJ, Kellis M. 2012. Efficient algorithms for the reconciliation problem with gene duplication, horizontal transfer and loss. Bioinformatics 28: 283–291.

Bartoli C, Lamichhane JR, Berge O, Guilbaud C, Varvaro L, Balestra GM, Vinatzer BA, Morris CE. 2015a. A framework to gauge the epidemic potential of plant pathogens in environmental reservoirs: The example of kiwifruit canker. Molecular Plant Pathology 16: 137–149.

Bartoli C, Carrere S, Lamichhane R, Varvaro L, Morris CE. 2015b. Whole-genome sequencing of 10 *Pseudomonas syringae* strains representing different host range spectra. Genome Announcements 3: 2–3.

Bender CL, Alarcón-Chaidez F, Gross DC. 1999. *Pseudomonas syringae* phytotoxins: Mode of action, regulation, and biosynthesis by peptide and polyketide synthetases. Microbiology and molecular biology reviews: MMBR 63: 266–292.

Berge O, Monteil CL, Bartoli C, Chandeysson C, Guilbaud C, Sands DC, Morris CE. 2014. A user’s guide to a data base of the diversity of *Pseudomonas syringae* and its application to classifying strains in this phylogenetic complex. PLoS ONE 9: e105547.

Berlin K, Koren S, Chin C-S, Drake JP, Landolin JM, Phillippy AM. 2015. Assembling large genomes with single-molecule sequencing and locality-sensitive hashing. Nature biotechnology 33: 623–630.

Bruns H, Crüsemann M, Letzel A-C, Alanjary M, McInerney JO, Jensen PR, Schulz S, Moore BS, Ziemert N. 2017. Function-related replacement of bacterial siderophore pathways. The ISME Journal: doi:10.1038/ismej.2017.

Buell CR, Joardar V, Lindeberg M, Selengut J, Paulsen IT, Gwinn ML, Dodson RJ, Deboy RT, Durkin AS, Kolonay JF, et al. 2003. The complete genome sequence of the *Arabidopsis* and tomato pathogen *Pseudomonas syringae* pv. *tomato* DC3000. Proceedings of the National Academy of Sciences of the United States of America 100: 10181–10186.

Bull CT, de Boer SH, Denny TP, Firrao G, Fischer-Le Saux M, Saddler GS, Scortichini M, Stead DE, Takikawa Y. 2010. Comprehensive list of names of plant pathogenic bacteria, 1980-2007. Journal of Plant Pathology 92: 551–592.

Bultreys A, Kaluzna M. 2010. Bacterial cankers caused by *Pseudomonas syringae* on stone fruit species with special emphasis on the pathovars *syringae* and *morsprunorum* Race 1 and Race 2. Journal of Plant Pathology 92: S1.21–S1.33.

Buonaurio R, Moretti C, da Silva DP, Cortese C, Ramos C, Venturi V. 2015. The olive knot disease as a model to study the role of interspecies bacterial communities in plant disease. Frontiers in plant science 6: 434.

Butler MI, Stockwell PA, Black MA, Day RC, Lamont IL, Poulter RTM. 2013. *Pseudomonas syringae* pv. *actinidiae* from recent outbreaks of kiwifruit bacterial canker belong to different clones that originated in China. PloS one 8: e57464.

Caballo-Ponce E, van Dillewijn P, Wittich R, Ramos C. 2016. WHOP, a genomic region associated with woody hosts in the *Pseudomonas syringae* complex contributes to the virulence and fitness of *Pseudomonas savastanoi* pv. *savastanoi* in olive plants. Molecular Plant-Microbe Interactions 30: 113–126.

Castresana J. 2000. Selection of conserved blocks from multiple alignments for their use in phylogenetic analysis. Molecular Biology and Evolution 17: 540–552.

Cohen O, Ashkenazy H, Belinky F, Huchon D, Pupko T. 2010. GLOOME: Gain loss mapping engine. Bioinformatics 26: 2914–2915.

Crosse, JE, & Garrett CME. 1966. Bacterial canker of stone-fruits. VII. Infection experiments with *Pseudomonas morsprunorum* and *Ps. syringae*. Annals of Applied Biology 58: 31–41.

Cunnac S, Chakravarthy S, Kvitko BH, Russell AB, Martin GB, Collmer A. 2011. Genetic disassembly and combinatorial reassembly identify a minimal functional repertoire of type III effectors in *Pseudomonas syringae*. Proceedings of the National Academy of Sciences of the United States of America 108: 2975–2980.

Darling AE, Mau B, Perna NT. 2010. Progressivemauve: Multiple genome alignment with gene gain, loss and rearrangement. PLoS ONE 5: e11147.

Dhillon BK, Laird MR, Shay JA, Winsor GL, Lo R, Nizam F, Pereira SK, Waglechner N, McArthur AG, Langille MGI, et al. 2015. IslandViewer 3: More flexible, interactive genomic island discovery, visualization and analysis. Nucleic Acids Research 43: W104–W108.

Ditta G, Stanfield S, Corbin D, Helinski DR. 1980. Broad host range DNA cloning system for gram-negative bacteria: construction of a gene bank of *Rhizobium meliloti*. Proceedings of the National Academy of Sciences of the United States of America 77: 7347–7351.

Dudnik A, Dudler R. 2014. Genomics-based exploration of virulence determinants and host-specific adaptations of *Pseudomonas syringae* strains isolated from grasses. Pathogens 3: 121–148.

Edwards DJ, Holt KE. 2013. Beginner’s guide to comparative bacterial genome analysis using next-generation sequence data. Microbial Informatics and Experimentation 3: 2.

Feil H, Feil WS, Chain P, Larimer F, DiBartolo G, Copeland A, Lykidis A, Trong S, Nolan M, Goltsman E, et al. 2005. Comparison of the complete genome sequences of *Pseudomonas syringae* pv. *syringae* B728a and pv. *tomato* DC3000. Proceedings of the National Academy of Sciences of the United States of America 102: 11064–11069.

Feil W, Feil H, Copeland A. 2012. Bacterial genomic DNA isolation using CTAB. URL http://1ofdmq2n8tc36m6i46scovo2e.wpengine.netdna-cdn.com/wp-content/uploads/2014/02/JGI-Bacterial-DNA-isolation-CTAB-Protocol-2012.pdf [accessed 28th December 2017]

Gardan L, Shafik H, Belouin S, Broch R, Grimont F, Grimont P. 1999. DNA relatedness among the pathovars of *Pseudomonas syringae* and description of *Pseudomonas tremae* sp. nov. and *Pseudornonas cannabina* sp. nov. (ex Sutic and Dowson 1959). International Journal of Systematic Bacteriology 49: 469–478.

Gardiner DM, Stiller J, Covarelli L, Lindeberg M, Shivas RG, Manners JM. 2013. Genome sequences of *Pseudomonas* spp. isolated from cereal crops. Genome announcements 1: e00209–e00213.

Gilbert V, Planchon V, Legros F, Maraite H, Bultreys A. 2009. Pathogenicity and aggressiveness in populations of *Pseudomonas syringae* from Belgian fruit orchards. European Journal of Plant Pathology 126: 263–277.

Grant M, Jones J. 2009. Hormone (dis)harmony moulds plant health and disease. Science 324: 750–752.

Green S, Studholme DJ, Laue BE, Dorati F, Lovell H, Arnold D, Cottrell JE, Bridgett S, Blaxter M, Huitema E, et al. 2010. Comparative genome analysis provides insights into the evolution and adaptation of *Pseudomonas syringae* pv. *aesculi* on *Aesculus hippocastanum*. PloS one 5: e10224.

Guttman DS, McHardy AC, Schulze-Lefert P. 2014. Microbial genome-enabled insights into plant–microorganism interactions. Nature Reviews Genetics 15: 797–813.

Hockett KL, Nishimura MT, Karlsrud E, Dougherty K, Baltrus DA. 2014. *Pseudomonas syringae* CC1557: A highly virulent strain with an unusually small type III effector repertoire That includes a novel effector. Molecular Plant-Microbe Interactions: MPMI 27: 923–32.

Hulin MT, Mansfield JW, Brain P, Xu X, Jackson RW, Harrison RJ. 2018. Characterisation of the pathogenicity of strains of *Pseudomonas syringae* towards cherry and plum. Plant Pathology. (Accepted article Dec 2017)

Hunt M, Silva N De, Otto TD, Parkhill J, Keane JA, Harris SR. 2015. Circlator: Automated circularization of genome assemblies using long sequencing reads. Genome Biology 16: 294.

Hurley B, Lee D, Mott A, Wilton M, Liu J, Liu YC, Angers S, Coaker G, Guttman DS, Desveaux D. 2014. The *Pseudomonas syringae* type III effector HopF2 suppresses *Arabidopsis* stomatal immunity. PLoS ONE 9: e114921.

Jackson RW, Athanassopoulos E, Tsiamis G, Mansfield JW, Sesma a, Arnold DL, Gibbon MJ, Murillo J, Taylor JD, Vivian a. 1999. Identification of a pathogenicity island, which contains genes for virulence and avirulence, on a large native plasmid in the bean pathogen *Pseudomonas syringae* pathovar *phaseolicola*. Proceedings of the National Academy of Sciences of the United States of America 96: 10875–80.

Joardar V, Lindeberg M, Jackson RW, Selengut J, Dodson R, Brinkac LM, Daugherty SC, Deboy R, Durkin AS, Giglio MG, et al. 2005. Whole-genome sequence analysis of *Pseudomonas syringae* pv. *phaseolicola* 1448A reveals divergence among pathovars in genes involved in virulence and transposition. Journal of Bacteriology 187: 6488–6498.

Jones JDG, Dangl JL. 2006. The plant immune system. Nature 444: 323–329.

Kamiunten H, Nakaol T, Oshida S. 2000. Agent of bacterial gall of cherry tree. J. Gen. Plant Pathol. 66: 219–224.

Katoh K, Misawa K, Kuma K, Miyata T. 2002. MAFFT: A novel method for rapid multiple sequence alignment based on fast Fourier transform. Nucleic acids research 30: 3059–3066.

Kearse M, Moir R, Wilson A, Stones-Havas S, Cheung M, Sturrock S, Buxton S, Cooper A, Markowitz S, Duran C, et al. 2012. Geneious Basic: An integrated and extendable desktop software platform for the organization and analysis of sequence data. Bioinformatics 28: 1647–1649.

Kim YJ, Lin NC, Martin GB. 2002. Two distinct *Pseudomonas* effector proteins interact with the Pto kinase and activate plant immunity. Cell 109: 589–598.

Kovach ME, Elzer PH, Hill DS, Robertson GT, Farris M a, Roop RM, Peterson KM. 1995. Four new derivatives of the broad-host-range cloning vector pBBR1MCS, carrying different antibiotic-resistance cassettes. Gene 166: 175–6.

Kvitko B, Collmer A. 2011. Construction of *Pseudomonas syringae* pv. *tomato* DC3000 mutant and polymutant strains. In: McDowell J, ed. Plant Immunity. Methods in Molecular Biology (Methods and Protocols). Humana Press, 109–128.

Kvitko BH, Park DH, Velásquez AC, Wei C-F, Russell AB, Martin GB, Schneider DJ, Collmer A. 2009. Deletions in the repertoire of *Pseudomonas syringae* pv. *tomato* DC3000 type III secretion effector genes reveal functional overlap among effectors. PLoS pathogens 5: e1000388.

Lamichhane JR, Varvaro L, Parisi L, Audergon J-M, Morris CE. 2014. Chapter Four - Disease and frost damage of woody plants caused by *Pseudomonas syringae:* Seeing the forest for the trees. In: Boyer J, Alexander M, Kamprath E, eds. Advances in Agronomy. San Diego: Academic Press, 235–295.

Langmead B, Salzberg S. 2013. Fast gapped-read alignment with Bowtie2. Nature methods 9: 357–359.

Larkin M, Blackshields G, Brown NP, Chenna R, Mcgettigan P a., McWilliam H, Valentin F, Wallace IM, Wilm A, Lopez R, et al. 2007. Clustal W and Clustal X version 2.0. Bioinformatics 23: 2947–2948.

Leong S, Ditta G, Helinshi D. 1982. Heme biosynthesis in *Rhizobium:* identification of a cloned gene coding for deltaaminolevulinic acid synthetase from *Rhizobium meliloti*. Journal of Biological Chemistry 257: 8724–8730.

Li H, Handsaker B, Wysoker A, Fennell T, Ruan J, Homer N, Marth G, Abecasis G, Durbin R. 2009. The Sequence Alignment/Map format and SAMtools. Bioinformatics 25: 2078–2079.

Li L, Stoeckert CJJ, Roos DS. 2003. OrthoMCL: Identification of ortholog groups for eukaryotic genomes. Genome Research 13: 2178–2189.

Lin NC, Martin GB. 2005. An *avrPto/avrPtoB* mutant of *Pseudomonas syringae* pv. *tomato* DC3000 does not elicit Pto-mediated resistance and is less virulent on tomato. Molecular Plant-Microbe Interactions MPMI 18: 43–51.

Lindgren PB. 1997. The role of *hrp* genes during plant-bacterial interactions. Annual review of phytopathology 35: 129–52.

Liu H, Qiu H, Zhao W, Cui Z, Ibrahim M, Jin G, Li B, Zhu B, Xie GL. 2012. Genome sequence of the plant pathogen *Pseudomonas syringae* pv. panici LMG 2367. Journal of Bacteriology 194: 5693–5694.

Liu W, Xie Y, Ma J, Luo X, Nie P, Zuo Z, Lahrmann U, Zhao Q, Zheng Y, Zhao Y, et al. 2015. IBS: An illustrator for the presentation and visualization of biological sequences. Bioinformatics 31: 3359–3361.

Lo T, Koulena N, Seto D, Guttman DS, Desveaux D. 2016. The HopF family of *Pseudomonas syringae* type III secreted effectors. Molecular Plant Pathology 18: 1–12.

Loman NJ, Quinlan AR. 2014. Poretools: A toolkit for analyzing nanopore sequence data. Bioinformatics 30: 3399–3401.

Mansfield J, Genin S, Magori S, Citovsky V, Sriariyanum M, Ronald P, Dow M, Verdier V, Beer SV, Machado MA, et al. 2012. Top 10 plant pathogenic bacteria in molecular plant pathology. Molecular Plant Pathology 13: 614–629.

Matas IM, Castañeda-Ojeda MP, Aragón IM, Antúnez-Lamas M, Murillo J, Rodriquez-Palenzuela P, Lopez-Solanilla E, Ramos C. 2014. Translocation and functional analysis of Pseudomonas savastanoi pv. savastanoi NCPPB 3335 type III secretion system effectors reveals two novel effector families of the *Pseudomonas syringae* complex. Molecular Plant-Microbe interactions: MPMI 27: 424–436.

Marcelletti S, Ferrante P, Petriccione M, Firrao G, Scortichini M. 2011. *Pseudomonas syringae* pv. *actinidiae* draft genomes comparison reveal strain-specific features involved in adaptation and virulence to *Actinidia* species. PLoS ONE 6: e27297.

Martínez-García PM, Rodríguez-Palenzuela P, Arrebola E, Carrión VJ, Gutièrrez-Barranquero JA, Pèrez-García A, Ramos C, Cazorla FM, De Vicente A. 2015. Bioinformatics analysis of the complete genome sequence of the mango tree pathogen *Pseudomonas syringae* pv. *syringae* UMAF0158 reveals traits relevant to virulence and epiphytic lifestyle. PLoS ONE 10: 1–26.

Mazzaglia A, Studholme DJ, Taratufolo MC, Cai R, Almeida NF, Goodman T, Guttman DS, Vinatzer BA, Balestra GM. 2012. *Pseudomonas syringae* pv. *actinidiae* (PSA) isolates from recent bacterial canker of kiwifruit outbreaks belong to the same genetic lineage. PLoS ONE 7: 1–11.

McCann HC, Rikkerink EHA, Bertels F, Fiers M, Lu A, Rees-George J, Andersen MT, Gleave AP, Haubold B, Wohlers MW, et al. 2013. Genomic analysis of the kiwifruit pathogen *Pseudomonas syringae* pv. *actinidiae* provides insight into the origins of an emergent plant disease. PLoS pathogens 9: e1003503.

McCann HC, Li L, Liu Y, Li D, Pan H, Zhong C, Rikkerink EHA, Templeton MD, Straub C, Colombi E, et al. 2017. Origin and evolution of the kiwifruit canker pandemic. Genome Biology and Evolution 9: 932–944.

Ménard M, Sutra L, Luisetti J, Prunier JP, Gardan L. 2003. *Pseudomonas syringae* pv. *avii* (pv. nov.), the causal agent of bacterial canker of wild cherries (*Prunus avium*) in France. European Journal of Plant Pathology 109: 565–576.

de Mendiburu F. 2016. Agricolae: Statistical procedures for agricultural research. URL https://cran.r-project.org/web/packages/agricolae/index.html [accessed 28th December 2017]

Monteil CL, Yahara K, Studholme DJ, Mageiros L, Méric G, Swingle B, Morris CE, Vinatzer BA, Sheppard SK. 2016. Population-genomic insights into emergence, crop-adaptation, and dissemination of *Pseudomonas syringae* pathogens. Microbial Genomics 21: e000089.

Moretti C, Cortese C, Passos da Silva D, Venturi V, Ramos C, Firrao G, Buonaurio R. 2014. Draft genome sequence of *Pseudomonas savastanoi* pv. *savastanoi* strain DAPP-PG 722, isolated in Italy from an olive plant affected by knot disease. Genome Announcements 2: e00864-14–e00864-14.

Mott GA, Thakur S, Smakowska E, Wang PW, Belkhadir Y, Desveaux D, Guttman DS. 2016. Genomic screens identify a new phytobacterial microbe-associated molecular pattern and the cognate *Arabidopsis* receptor-like kinase that mediates its immune elicitation. Genome Biology 17: 98.

Moulton J, Vivian A, Hunter P, Taylor JD. 1993. Changes in cultivar-specificity toward pea can result from transfer of plasmid RP4 and other incompatibility group P1 replicons to *Pseudomonas syringae* pv. *pisi*. Journal of General Microbiology 39: 3149–3155.

Nahar K, Matsumoto I, Taguchi F, Inagaki Y, Yamamoto M, Toyoda K, Shiraishi T, Ichinose Y, Mukaihara T. 2014. *Ralstonia solanacearum* type III secretion system effector Rip36 induces a hypersensitive response in the nonhost wild eggplant *Solanum torvum*. Molecular Plant Pathology 15: 297–303.

Neale HC, Laister R, Payne J, Preston G, Jackson RW, Arnold DL. 2016. A low frequency persistent reservoir of a genomic island in a pathogen population ensures island survival and improves pathogen fitness in a susceptible host. Environmental Microbiology 18: 4144–4152.

Neale HC, Slater RT, Mayne L-M, Manoharan B, Arnold DL. 2013. In planta induced changes in the native plasmid profile of *Pseudomonas syringae* pathover *phaseolicola* strain 1302A. Plasmid 70: 420–424.

Nowell RW, Laue BE, Sharp PM, Green S. 2016. Comparative genomics reveals genes significantly associated with woody hosts in the plant pathogen *Pseudomonas syringae*. Molecular Plant Pathology 17: 1409–1424.

O’Brien HE, Thakur S, Gong Y, Fung P, Zhang J, Yuan L, Wang PW, Yong C, Scortichini M, Guttman DS. 2012. Extensive remodeling of the *Pseudomonas syringae* pv. *avellanae* type III secretome associated with two independent host shifts onto hazelnut. BMC microbiology 12: 141.

O’Brien HE, Thakur S, Guttman DS. 2011. Evolution of plant pathogenesis in *Pseudomonas syringae:* a genomics perspective. Annual review of phytopathology 49: 269–289.

Pagel M. 2004. Detecting correlated evolution on phylogenies: A general method for the comparative analysis of discrete characters. Proceedings of the Royal Society of London. Series B, Biological Sciences 255: 37–45.

Paradis E, Claude J, Strimmer K. 2004. APE: Analyses of phylogenetics and evolution in R language. Bioinformatics 20: 289–290.

Parkinson N, Bryant R, Bew J, Elphinstone J. 2011. Rapid phylogenetic identification of members of the *Pseudomonas syringae* species complex using the *rpoD* locus. Plant Pathology 60: 338–344.

Pitman AR, Jackson RW, Mansfield JW, Kaitell V, Thwaites R, Arnold DL. 2005. Exposure to host resistance mechanisms drives evolution of bacterial virulence in plants. Current Biology: CB 15: 2230–2235.

Posada D. 2008. jModelTest: Phylogenetic model averaging. Molecular Biology and Evolution 25: 1253–1256.

Press MO, Li H, Creanza N, Kramer G, Queitsch C, Sourjik V, Borenstein E. 2013. Genome-scale co-evolutionary inference identifies functions and clients of bacterial Hsp90. PLoS Genetics 9: e1003631.

Qi M, Wang D, Bradley CA, Zhao Y. 2011. Genome sequence analyses of *Pseudomonas savastanoi* pv. *glycinea* and subtractive hybridization-based comparative genomics with nine pseudomonads. PLoS ONE 6: e16451.

Quevillon E, Silventoinen V, Pillai S, Harte N, Mulder N, Apweiler R, Lopez R. 2005. InterProScan: Protein domains identifier. Nucleic Acids Research 33: 116–120.

R Core Team. 2012. R: A language and environment for statistical computing. Vienna, Austria: R Foundation for Statistical Computing.

Ravindran A, Jalan N, Yuan JS, Wang N, Gross DC. 2015. Comparative genomics of *Pseudomonas syringae* pv. *syringae* strains B301D and HS191 and insights into intrapathovar traits associated with plant pathogenesis. Microbiology Open 4: 553–573.

Rezaei R, Taghavi SM. 2014. Host specificity, pathogenicity and the presence of virulence genes in Iranian strains of *Pseudomonas syringae* pv. *syringae* from different hosts. Archives of Phytopathology and Plant Protection 47: 2377–2391.

Rodríguez-Palenzuela P, Matas IM, Murillo J, López-Solanilla E, Bardaji L, Pérez-Martínez I, Rodríguez-Moskera ME, Penyalver R, López MM, Quesada JM, et al. 2010. Annotation and overview of the *Pseudomonas savastanoi* pv. *savastanoi* NCPPB 3335 draft genome reveals the virulence gene complement of a tumour-inducing pathogen of woody hosts. Environmental microbiology 12: 1604–1620.

Şahin F. 2001. Severe outbreak of bacterial speck, caused by *Pseudomonas syringae* pv. *tomato*, on field-grown tomatoes in the eastern Anatolia region of Turkey. Plant Pathology 50: 799.

Sarkar SF, Gordon JS, Martin GB, Guttman DS. 2006. Comparative genomics of host-specific virulence in *Pseudomonas syringae*. Genetics 174: 1041–1056.

Sawada H, Suzuki F, Matsuda I, Saitou N. 1999. Phylogenetic analysis of *Pseudomonas syringae* pathovars suggests the horizontal gene transfer of argK and the evolutionary stability of *hrp* gene cluster. Journal of molecular evolution 49: 627–644.

Sawada H, Shimizu S, Miyoshi T, Shinozaki T, Kusumoto S, Noguchi M, Naridomi T, Kikuhara K, Kansako M, Fujikawa T, et al. 2015. *Pseudomonas syringae* pv. *actinidiae* biovar 3. Japanese Journal of Phytopathology 81: 111–126.

Sawyer S. 1989. Statistical tests for detecting gene conversion. Molecular Biology and Evolution 6: 526–538.

Scholz-Schroeder BK, Hutchison ML, Grgurina I, Gross DC. 2001. The contribution of syringopeptin and syringomycin to virulence of *Pseudomonas syringae* pv. *syringae* strain B301D on the basis of *sypA* and *syrB1* biosynthesis mutant analysis. Molecular Plant-Microbe Interactions: MPMI 14: 336–348.

Schulze-Lefert P, Panstruga R. 2011. A molecular evolutionary concept connecting nonhost resistance, pathogen host range, and pathogen speciation. Trends in plant science 16: 117–125.

Scortichini M. 2010. Epidemiology and predisposing factors of some major bacterial diseases of stone and nut fruit trees species. Journal of Plant Pathology 92: 73–78.

Shafer A, Tauch A, Jager W, Kalinowski J, Thierbach G, Puhler A. 1994. Small mobilizable multi-purpose cloning vectors derived from the *Escherichia coli* plasmids pK18 and pK19: selection of defined deletions in the chromosome of *Corynebacterium glutamicum*. Gene 145: 69–73.

Sohn KH, Zhang Y, Jones JDG. 2009. The *Pseudomonas syringae* effector protein, AvrRPS4, requires in planta processing and the KRVY domain to function. Plant Journal 57: 1079–1091.

Stamatakis A. 2014. RAxML version 8: A tool for phylogenetic analysis and post-analysis of large phylogenies. Bioinformatics 30: 1312–1313.

Staskawicz BJ, Dahlbeck D, Keen NT. 1984. Cloned avirulence gene of *Pseudomonas syringae* pv. *glycinea* determines race-specific incompatibility on *Glycine max* (L.) Merr. Proceedings of the National Academy of Sciences of the United States of America 81: 6024–6028.

Thakur S, Weir BS, Guttman D. 2016. Phytopathogen genome announcement: Draft genome sequences of 62 *Pseudomonas syringae* type and pathotype strains. Molecular Plant-Microbe Interactions 29: 243–246.

Vicente JG, Alves JP, Russell K, Roberts SJ. 2004. Identification and discrimination of *Pseudomonas syringae* isolates from wild cherry in England. European Journal of Plant Pathology 110: 337–351.

Visnovsky SB, Fiers M, Lu A, Panda P, Taylor R, Pitman AR. 2016. Draft genome sequences of 18 strains of *Pseudomonas* isolated from kiwifruit plants in New Zealand and overseas. Genome Announcements 4: e00061–e00016.

Walker BJ, Abeel T, Shea T, Priest M, Abouelliel A, Sakthikumar S, Cuomo CA, Zeng Q, Wortman J, Young SK, et al. 2014. Pilon: An integrated tool for comprehensive microbial variant detection and genome assembly improvement. PLoS ONE 9: e112963.

Wang Y, Li J, Hou S, Wang X, Li Y, Ren D, Chen S, Tang X, Zhou J-M. 2010. A *Pseudomonas syringae* ADP-ribosyltransferase inhibits Arabidopsis mitogen-activated protein kinase kinases. The Plant cell 22: 2033–2044.

Warnes G, Bolker B, Bonebakker L, Gentleman R, Huber W, Liaw A, Lumley T, Maechler M, Magnusson A, Moeller S, et al. 2016. gplots: Various R programming tools for plotting data. URL https://cran.r-project.org/web/packages/gplots/index.html [accessed 28th December 2017]

Wu S, Lu D, Kabbage M, Wei H-L, Swingle B, Records AR, Dickman M, He P, Shan L. 2011. Bacterial effector HopF2 suppresses *Arabidopsis* innate immunity at the plasma membrane. Molecular Plant-Microbe Interactions: MPMI 24: 585–593.

Xiang T, Zong N, Zou Y, Wu Y, Zhang J, Xing W, Li Y, Tang X, Zhu L, Chai J, et al. 2008. *Pseudomonas syringae* effector AvrPto blocks innate immunity by targeting receptor kinases. Current Biology: CB 18: 74–80.

Young J. 1991. Pathogenicity and identification of the lilac pathogen, *Pseudomonas syringae* pv. *syringae* van Hall 1902. Ann. Appl. Biol. 118: 283–298.

Young JM. 2010. Taxonomy of *Pseudomonas syringae*. Journal of Plant Pathology 92: S1.5–S1.14.

Yu D, Yin Z, Li B, Jin Y, Ren H, Zhou J, Zhou W, Liang L, Yue J, Xu S. 2016. Gene flow, recombination, and positive selection in *Stenotrophomonas maltophilia:* Mechanisms underlying the diversity of the widespread opportunistic pathogen. Genome 59: 1063–1075.

Yu G, Smith D, Zhu H, Guan Y, Lam T. 2017. ggtree: An R package for visualization and annotation of phylogenetic trees with their covariates and other associated data. Methods in Ecology and Evolution 8: 28–36.

Zhang J, Li W, Xiang T, Liu Z, Laluk K, Ding X, Zou Y, Gao M, Zhang X, Chen S, et al. 2010. Receptor-like cytoplasmic kinases integrate signaling from multiple plant immune receptors and are targeted by a *Pseudomonas syringae* effector. Cell host & microbe 7: 290–301.

Zhao Y, Ma Z, Sundin GW. 2005. Comparative genomic analysis of the pPT23A plasmid family of *Pseudomonas syringae*. Journal of Bacteriology 187: 2113–2126.

Zhao W, Jiang H, Tian Q, Hu J. 2015. Draft genome sequence of *Pseudomonas syringae* pv. *persicae* NCPPB 2254. 3: 54–55.

